# *Drosophila* Toll-2 controls homeotic domain formation and timing of folding

**DOI:** 10.1101/2025.01.07.631521

**Authors:** Lale Alpar, Rémi Pigache, Adrien Leroy, Mehdi Ech-Chouini, Stéphane Pelletier, Yohanns Bellaïche

## Abstract

Tissue compartmentalization is a fundamental feature of animal development, crucial for determining cell fate, growth, and organ morphogenesis. Hox genes play essential and conserved roles in specifying homeotic compartments along the antero-posterior axis of the body. Despite extensive characterization of homeotic gene regulation, the mechanisms underlying the formation, maintenance, and reshaping of homeotic compartments during development remain incompletely understood. Here, we investigate the role of the Toll-2 receptor in the formation and folding of the Deformed (Dfd) homeotic compartment, which produces the neck of adult *Drosophila*. We show that Toll-2 is essential for properly forming the bilateral Dfd compartment through the fusion of the two eye-antennal imaginal discs. Additionally, we find that Toll-2 regulates the timing of neck morphogenesis by promoting cell apoptosis and cell-cell rearrangements preceding neck folding. Last, we establish that Toll-2 independently and sequentially governs both homeotic compartment formation and the timing of its morphogenesis via PI3K and Src signaling, respectively. Altogether, our findings provide novel insights into the spatiotemporal regulation of homeotic compartment development.

## Introduction

Hox genes encode a conserved family of transcription factors that regionalize the bilaterally symmetric body plan along its anterior-posterior (a-p) axis (Hubert and Wellik, 2024). The spatiotemporal patterns of Hox gene expression, as well as their critical roles in defining bilateral homeotic compartments, are well-documented. However, understanding the mechanisms by which Hox gene compartments are formed, maintained, and reshaped during development remains a central focus of research in development across the animal kingdom.

The function of homeotic genes was first discovered in *Drosophila* and later identified to be conserved in a wide range of species including mammals (Hubert and Wellik, 2024; Lewis, 1978; Nüsslein-Volhard and Wieschaus, 1980). In *Drosophila* and mammalian embryos, Hox genes are expressed in bilateral compartments along the a-p axis, specifying the cellular repertoires, size, and shape of each compartment (Hubert and Wellik, 2024). Such a-p expression is also found in adults, both in flies and mammals, despite the extensive and complex changes in morphology that occurs during development. The complexity of this morphogenetic process is particularly evident in *Drosophila*, wherein the bilaterally symmetrical imaginal discs need to be fused along the midline and profoundly remodeled during metamorphosis to give the adult head, thorax, and appendage structures. Mechanisms promoting dorsal thorax disc fusion —involving signaling pathways such as JNK and Fat/Dachsous— have been the focus of various studies (Athilingam et al., 2022; Lu et al., 2022; Martín-Blanco et al., 2000; Usui and Simpson, 2000; Zeitlinger and Bohmann, 1999). Yet, little attention has been given to understanding how homeotic compartments are fused and maintained during the complex morphogenetic movements that reshape larval imaginal discs into the prospective adult epithelium. More broadly, across the animal kingdom, seminal studies have delineated how Hox gene patterns are specified during a-p segmentation. Yet, our understanding of the *de novo* formation and maintenance of Hox gene compartments during subsequent morphogenesis remains limited.

*Deformed (Dfd)* is a Hox gene whose functional characterization has been instrumental in delineating how Hox-regulated transcriptional programs specify structures along the a-p axis (Lohmann et al., 2002; Stultz et al., 2012; Zhai et al., 2010). Dfd is critical for the development of both embryonic and adult *Drosophila* head structures (Anais Tiberghien et al., 2015; Diederich et al., 1991; Lohmann et al., 2002; Merrill et al., 1987). Previously, we identified a central role for Dfd in forming the *Drosophila* neck (Villedieu et al., 2023). In the *Drosophila* pupal epithelium, *Dfd* is expressed in a narrow a-p compartment at the interface between the head and thorax, corresponding to the prospective neck of the adult. Within this compartment, Dfd regulates actomyosin enrichment and tension generation via Toll-like receptor (TLR) Toll-8 in order to enhance the rate of tissue deepening during neck folding. The TLR family of cell surface receptors is a conserved group of molecules best known for their roles in immune response and embryonic patterning, although a growing body of work highlights their roles in controlling cell dynamics with morphogenetic outcomes (Iijima et al., 2020; Lavalou et al., 2021; Paré et al., 2019, 2014). Interestingly, the expression of cell surface receptors, such as TLRs can contribute to tissue compartment identity downstream of selector genes (Fischer et al., 2024; Hsia et al., 2017; Iijima et al., 2020; Milán et al., 2001; Tepass et al., 2002; Wang and Dahmann, 2020). Relevantly, a subset of TLRs are expressed in striped patterns along the a-p axis and control germ band elongation in the early *Drosophila* embryo (Paré et al., 2014). However, whether and how TLRs contribute to shaping and maintaining homeotic compartments during morphogenesis remains unexplored.

Here, we aim to investigate how the pupal *Dfd* homeotic compartment forms during metamorphosis and how its subsequent morphogenesis is controlled. Using a combination of live imaging and fixed tissue analyses, we delineated the complex 3D morphogenetic choreography of Dfd compartment fusion prior to head eversion. By screening for cell surface receptors, we identified a specific role for Toll-2 in controlling the formation of an a-p band of neck cells and provided evidence that Toll-2 mediates this role through regulation of actomyosin dynamics, which are necessary for the correct sorting of *Dfd-*positive (*Dfd+*) cells. Furthermore, we found that within the *Dfd* homeotic compartment, Toll-2 regulates both cell delamination and rearrangement rates, influencing the onset of neck folding without impacting the rate of tissue deepening during folding. Together, our work advances the understanding of mechanisms underlying homeotic compartment formation and morphogenesis during development.

## Results

### *Dfd* neck compartment originates from an anterior subset of cells in the eye-antennal *imaginal* discs

To construct the bilaterally symmetrical and continuous body plan, the left and right imaginal discs of the head, thorax and appendages need to fuse with each other, and with the neighboring tissues during metamorphosis (Milner et al., 1984; Usui and Simpson, 2000). Following disc fusion, the head, initially enclosed within the thoracic segment, everts around 12 hours after pupa formation (hAPF), bringing the pupal neck to its final position along the a-p axis, at the interface with the neighboring dorsal thorax (Bainbridge and Bownes, 1981; Villedieu et al., 2023). Having found that the dorsal part of the adult neck is marked by *Dfd* expression (Villedieu et al., 2023), we aimed to trace back the *Dfd+* precursors of the adult *Drosophila* neck, to their origins in the earlier developmental stages. Hence, we followed *Dfd+* cells in the larval precursors of the dorsal head and thorax, the eye-antennal and wing imaginal discs respectively. This was achieved by using a Gal4 knock-in at the *Dfd* locus driving the expression of *UAS-LifeAct::Ruby (Dfd>LifeAct::Ruby*), thereafter used as a proxy for *Dfd* expression. In 3^rd^ instar larvae, *Dfd>LifeAct::Ruby* signal was detected in the eye-antennal but not in the wing imaginal disc (Figure 1A, Figure S1A, B). At this stage, the left and right eye-antennal discs have not yet fused, though a thin cellular membrane continuous to and joining the two is present (Figure S1B, Figure 2H) (Haynie and Bryant, 1986; Milner et al., 1984). *Dfd>LifeAct::Ruby* was expressed in both eye discs, flanking but not reaching across this connection (Figure S1B). Notably, *Dfd>LifeAct::Ruby* expression was restricted to the peripodial layer of the disc, starting from the most anterior tips, where the discs were attached to the cephalopharyngeal skeleton (Haynie and Bryant, 1986) and extending slightly further than the boundary of the eye and antennal portions of the disc (Figure 1A, Figure S1B). Strikingly, and as established before (Haynie and Bryant, 1986), the peripodial layer of the eye-antennal disc at this anterior-most level was columnar rather than squamous, equal in thickness to the underlying disc proper (Figure 1A). More importantly, these columnar peripodial cells were reported to contribute to the dorsal and posterior sections of the adult head (Haynie and Bryant, 1986), including the structures that show defects when Dfd function is lost (Merrill et al., 1987; Villedieu et al., 2023). This is consistent with the possibility that the *Dfd+* neck compartment originates from a subset of peripodial cells, present in both left and right eye-antennal discs, that fuse together during the early stages of pre-pupal development to form the pupal *Dfd+* neck compartment.

**Figure 1:**
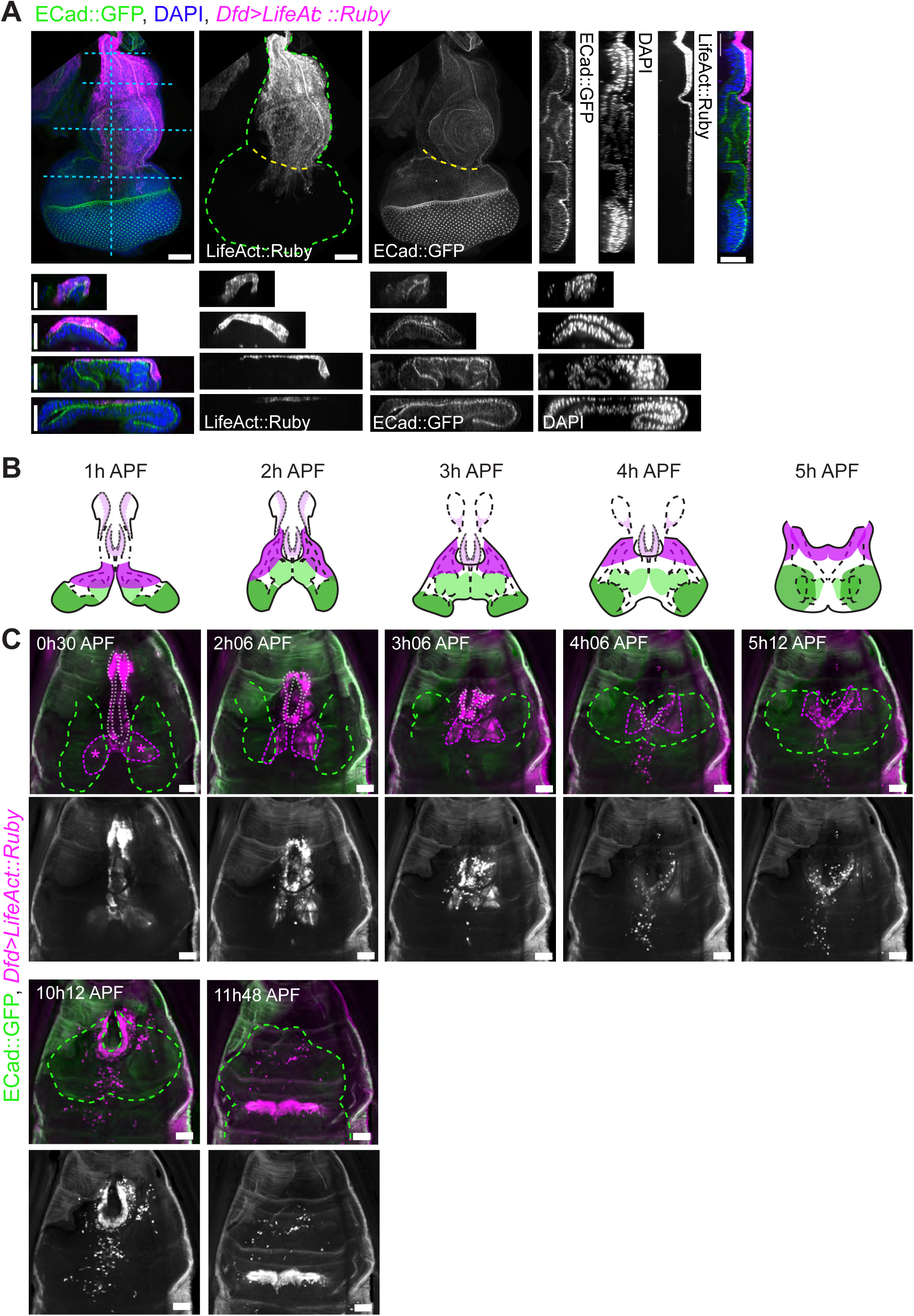
Dfd expression in the larvae and the early pupa. A: 3^rd^ instar eye-antennal discs expressing *UAS*-*LifeAct::Ruby* under *DfdGal4* control. Dashed blue lines mark position of reslices: (right) anterior-posterior reslice, (below) series of medial-lateral relices moving from anterior to posterior, ordered top to bottom. Scale bars = 50 μm. B: Schematics summarizing the changes in *Dfd*+ compartment position within eye-antennal discs over time, based on the images shown in C, Movie 1 and Sup Fig 1C. Dark green regions = eye, light green regions = antenna, magenta regions = *Dfd* expression (expression outside the eye-antennal discs is colored lighter). C: Snapshots taken from Movie 1, showing position of *DfdGal4>UAS-LifeAct::Ruby* expressing tissue within the early pupa. Green dashed lines outline position of wing imaginal discs/nota, white dashed lines outline *Dfd* expression outside the eye-antennal discs, magenta dashed lines outline *Dfd* expression within eye-antennal discs. Initial position of *Dfd+* eye-antennal compartments is marked with asterisks in first panel. Scale bars = 100 μm.

**Figure 2:**
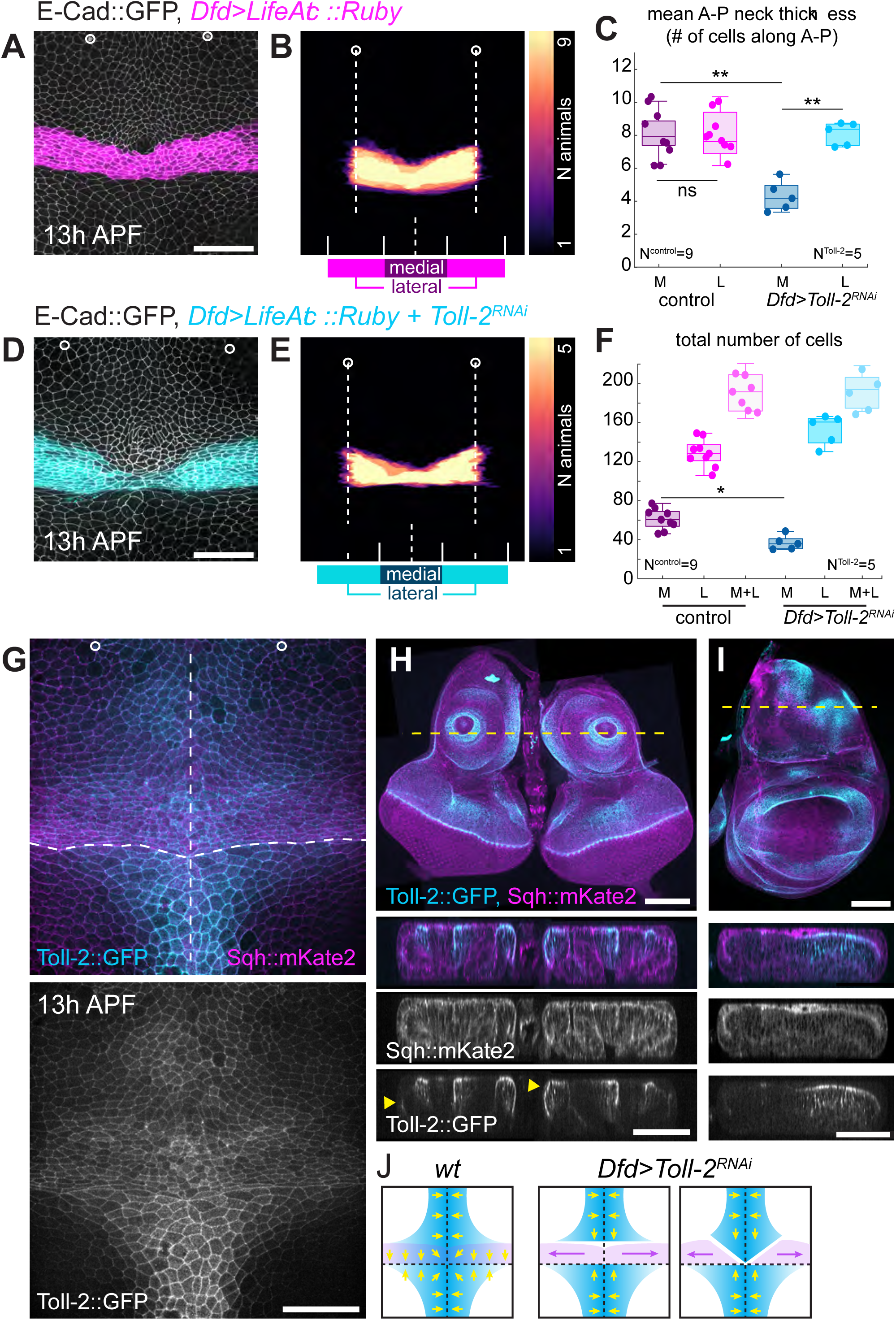
Loss of Toll-2 function in the *Dfd+* compartment results in defective compartment shape. A: *DfdGal4>UAS-LifeAct::Ruby* and E-Cad::GFP pupa at 13 hAPF illustrating the shape of the *Dfd+* neck compartment in the dorsal-medial region of the pupa. Scale bars: 50 µm. B: *DfdGal4>UAS-LifeAct::Ruby* expressing region between head macrochaetes for different animals (at 14h APF) segmented and overlayed. White circles and long dashed lines mark positions of head macrochaetes used as landmarks. Bottom of the panel: short dashed line marks position of midline. Solid white lines mark limits of the medial (dark purple) and lateral (magenta) regions defined for quantifications shown in C, and F, corresponding to the medial half of the region between macrochaetes and the two laterally adjacent regions of the same size, respectively. C: Average number of cells (mean ± sem) found along the anterior-to-posterior (a-p) direction in medial (dark shades) and lateral (light shades) neck regions of) *DfdGal4>UAS-LifeAct::Ruby* expressing control (magenta) or *DfdGal4>UAS-LifeAct::Ruby* + *UAS-Toll-2^RNAi^* (blue) animals, averaged over 13 to 15 h APF. Box plots represent the median and the upper quartiles of data plotted. Each data point represents average vale for one animal. N(control) = 9 pupae, N(*Toll-2^RNAi^*) = 5 pupae. ** p-value < 0.01, Welch test. D: Neck shape as described by *DfdGal4>UAS-LifeAct::Ruby* expression in the neck region of a *DfdGal4>UAS-LifeAct::Ruby* + *UAS-Toll-2^RNAi^* animal at 13 h APF. Scale bars = 50 µm. E: *DfdGal4>UAS-LifeAct::Ruby* expressing region between head macrochaetes for different *DfdGal4> UAS-LifeAct::Ruby* + *UAS-Toll-2^RNAi^* animals (at 14h APF) segmented and overlayed. White circles and long dashed lines mark positions of head macrochaetes used as landmarks. Bottom of the panel: short dashed line marks position of midline. Solid white lines mark limits of the medial (dark blue) and lateral (cyan) regions defined for quantifications shown in C and F, corresponding to the medial half of the region between macrochaetes and the two laterally adjacent regions of the same size, respectively. F: Total number of cells allocated to each region (medial = dark, lateral = medium, or total of both = light shades), in *DfdGal4>UAS-LifeAct::Ruby* control (magenta) and *DfdGal4> UAS-LifeAct::Ruby* + *UAS-Toll-2^RNAi^* (blue) animals, averaged over 13 to 15 h APF. Box plots represent the median and the upper quartiles of data plotted. Each data point represents average vale for one animal. N(control) = 9 pupae, N(*Toll-2^RNAi^*) = 5 pupae. * p-value < 0.05, Welch test. G: Toll-2::GFP and Sqh::mKate2 in 13h APF pupa. Circles mark head macrochaete positions. Dashed lines mark midline and head/thorax boundary (i.e. where the left/right and eye/wing discs meet). Scale bar = 50 μm. H: Toll-2::GFP and Sqh::mKate2 in eye-antennal imaginal discs. Lower panels show resliced sections taken along the apical-basal axis, at the position marked by yellow dashed line. Note the inter-antennal connection linking the two discs. Arrowheads mark where Toll-2 can be seen in the peripodia. Scale bars = 50 μm. C: Toll-2::GFP and Sqh::mKate2 in the wing imaginal disc. Lower panels show resliced sections taken along the apical-basal axis, at the position marked by yellow dashed line. Scale bars = 50 μm. D: Schematics showing potential explanation for *Dfd>Toll-2^RNAi^* neck shape phenotype. In the wildtype animals, Toll-2 (blue) is found in a cross-like pattern at the intersection of four discs, hypothesized to help cells at the interface to find and fuse with each other (yellow arrows). When Toll-2 is specifically removed from the neck region, the left and right *Dfd+* neck cells (magenta) will have diminished capacity to meet, while the medial head and notum cells will still be able to fuse. This may bring the head and notum midlines together at the expense of the medial *Dfd+* cells that get pushed towards the lateral.

To follow the *Dfd+* neck compartment precursors during pre-pupal development stages, we performed time-lapse imaging of early pupal development from 0 hAPF to head eversion. As pupae do not survive the removal of the pupal case prior to head eversion, we imaged *Dfd>LifeAct::Ruby* with E-Cad::GFP through the pupal case (movie 1, Figure 1B, C). To supplement these movies, we also dissected and fixed pre-pupal tissues at various early stages, which allowed for confocal imaging at higher resolution and better signal-to-noise ratio. Matching the *Dfd>LifeAct::Ruby* regions in these fixed-tissue images and time-lapse movies, we were able to identify and follow the relative position of various *Dfd+* anatomical structures throughout early pupal stages (Figure 1B, C, Figure S1C). At 1 hAPF, *Dfd>LifeAct::Ruby* expression was detected in the labial discs and in the cells surrounding the cibarium, in addition to the eye-antennal discs, where the pattern was reminiscent of that seen in the larva. At this stage, *Dfd>LifeAct::Ruby* anterior peripodia of each disc could be identified flanking the two sides of the strongly marked cibarium (Figure 1C and Figure S1C, first panels). Images of fixed tissues and their 3D reconstruction revealed that left and right eye-antennal discs had already started their fusion as early as 2 hAPF. However, the two *Dfd+* compartments were not yet in contact at these early stages (Figure S1C, D, panels 1-3). From 1 to 4 hAPF, the eye-antennal discs continued to fuse progressively, while the left and right *Dfd+* compartments appeared to rotate towards the midline. At around 5 hAPF, the left and right *Dfd+* compartments medially approached each other (Figure S1C, bottom panel). At the same time, an overall compaction and movement in the anterior direction gave an obtuse triangular shape to each compartment (Figure S1C, bottom panels). To complement these images and get a better understanding of tissue architecture at this point, we also imaged whole pupa at 5 hAPF by microCT scanning and segmented the eye-antennal discs. These 3D reconstructions and segmentations revealed a 3D geometry confirming the arrangements of the eye-antennal disc deduced from fixed and live confocal microscopy (Movie 2).

Using the fixed images and their 3D reconstructions to guide us, we could observe a sequence of events in time-lapse images of live pre-pupa (Figure 1C, Movie 1, schematically summarized in Figure 1B). Following the *Dfd>LifeAct::Ruby* eye-antennal tissues from their initial position *in vivo*, we found that they moved anteriorly and then compacted along the anterior-posterior axis within the first 4 hours (Movie 1; Figure 1C, panels 1-4). At around 3-4h APF, we observed the beginning of the eversion and displacements of the wing disc nota towards each other, while the eye-antennal discs rotated ventrolaterally (Movie 1). Presumably due to these movements and the subsequent fusion of the nota, live detection of the eye-antennal *Dfd>LifeAct::Ruby* signal became obscure for a brief period around 4 hAPF. The *Dfd>LifeAct::Ruby* disc compartments were then clearly visible from around 5h APF, at which point, the triangular left and right compartments in contact collectively appeared as a V-shape. This V-shape gradually evolved into a U-shaped band by 6 hAPF and eventually expanded during head eversion at 12h APF to form the dorsally located *Dfd+* neck compartment (Movie 1, Figure 1C, bottom panels). Collectively, we concluded that the *Dfd+* compartments residing in the anterior tips of the left and right eye-antennal discs of the 3^rd^ instar larva gradually moves to meet each other medially within the first 5 hours of the pupal development to form the bilateral symmetric *Dfd+* pupal neck compartment.

### Toll-2 is required for bringing together the two halves of the *Dfd+* neck compartment

Developmental compartments are often associated with specific cell-cell adhesion properties that promote cohesion among cells that express the same selector gene, and prevent them from mixing across compartment boundaries (Mann, 2000; Tepass et al., 2002; Wang and Dahmann, 2020). We hypothesized that cell-cell adhesion may help the left and right *Dfd+* neck compartments to adhere to each other to form the bilateral symmetric compartment.

Following head eversion, the *Dfd+* neck compartment was an unbroken band that spans the medial-lateral (m-l) body axis (Figure 2A). The a-p thickness of this band was close to uniform along the m-l axis, and showed little variation across animals (Figure 2A-C). Assuming that failure to fuse the left and right *Dfd+* neck compartments would disrupt this uniform shape, we checked, as a proof of principle, if loss of adhesion can impact neck shape, by reducing E-Cadherin (E-Cad, Shotgun, shg, in *Drosophila*) level within the Dfd domain (*Dfd>shg^RNAi^*). In these animals, the a-p thickness of the *Dfd*-expressing band of cells was unusually thinner in the medial-most part of the neck where the left and right halves were expected to meet (Figure S2) – in the most extreme case, the *Dfd+* cells failed to meet in the medial domain of the neck; thus causing the head and thorax epithelia to be in direct contact. This suggests that cell-cell adhesion within the *Dfd+* compartment is necessary to control the fusion of the left and right haves of the *Dfd+* neck compartment. We therefore aimed to determine whether other cell-cell adhesion molecules would regulate the shape of the compartment.

Previously, we have identified the cell surface receptor Toll-8, to be enriched specifically in the Dfd compartment, and to regulate its folding (Villedieu et al., 2023). While the absence of Toll-8 itself did not impact the initial uniform shape of the *Dfd+* compartment (Figure S2 and (Villedieu et al., 2023), we wondered if other surface adhesion proteins of the same family would mirror the loss of E-Cad function. Strikingly, knocking down *Toll-2* in the *neck* (*Dfd>Toll-2^RNAi^*) significantly reduced the number of cell rows in the medial region of the neck, resulting in a drastically lower a-p neck thickness as observed in *Dfd>shg^RNAi^* animals (Figure 2C-E). Though neck shape was variable in *Dfd>Toll-2^RNAi^* animals (Figure 2E), this lower a-p thickness in the most medial neck domain was consistent across animals (Figure 2C and E). Expanding our search to other Toll-like receptors, and members of the larger leucine-rich repeat (LRR) domain superfamily that they belong to, did not yield other genes whose knock-down impact the initial *Dfd+* compartment shape (Figure S2). We concluded that Toll-2 is required in the *Dfd+* neck compartment for shaping this compartment formed by the fusion of the two eye-antennal imaginal discs.

### Toll-2 maintains together *Dfd+* cells during disc fusion

To better understand how Toll-2 control the shape of the *Dfd+* neck compartment in the pre-pupa, we analyzed the pattern of *Dfd>LifeAct::Ruby* in *Dfd>Toll-2^RNAi^* tissue from the 3^rd^ instar larval stage onwards. While we could not observe striking differences prior to eye-antennal disc fusion, we found that once the bilateral *Dfd+* domain becomes visible as a V-shaped band around 5 h APF, defects in the shape of this band, such as narrower or discontinuous medial connections, can be readily observed in *Dfd>Toll-2^RNAi^* animals (Figure S3). To understand this phenotype better we generated a Toll-2::GFP knock-in to characterize its protein localization in control animals. In the pupa, Toll-2::GFP is present in the neck as well as in the neighboring head and notum midline regions (Figure 2G). Accordingly, Toll-2::GFP can be found in both eye-antennal as well as wing imaginal discs in the 3^rd^ instar larvae (Figure 2H, I). Consistently with its role in the neck compartment, Toll-2::GFP was not restricted to the disc epithelium proper. Furthermore, Toll-2::GFP could be observed in the medial edges of the discs, near the inter-antennal connection (Figure 2G, lower panels). In a similar manner, Toll-2::GFP in the pupa also located to the parts of the head and notum where the two body halves meet. Indeed, Toll-2::GFP pattern in the pupa was reminiscent of a cross, marking the regions contributing to the fusion of the four imaginal discs (i.e. left and right eye-antennal and wing imaginal discs) to make up the dorsal head and thorax epithelium (Figure 2G and J). Building on this characterization of Toll-2:GFP localization, we aimed to understand how Toll-2 expression within the *Dfd+* compartment contributed to neck shape.

Given Toll-2’s known role as a surface adhesion receptor (Paré et al., 2014), we hypothesized that Toll-2 might be important for the cells at the limits of these discs to adhere to each other. If this hypothesis was correct, removing Toll-2 specifically from the *Dfd+* cells would disrupt the fusion of the left and right halves of this domain without affecting the total number of cells within the Dfd+ compartment. Indeed, in *Dfd>Toll-2^RNAi^* animals, the total number of medial neck cells was reduced, alongside a slight increase in the number of lateral cells (Figure 2F). This agreed with a possible cell sorting defect that brings together the medial head and notum cells at the expense of displacing the medial neck cells towards the lateral (Figure 2J). To further explore the possibility of a cell sorting function of Toll-2, we analyzed the distribution of *Toll-2^RNAi^* clones within control tissues. When surrounded by control cells, neck *Toll-2^RNAi^* clones were characterized by a smooth interface, marked by an enrichment of Myosin II (Myo II) (Figure 3A). The *Toll-2^RNAi^* clone roundness and MyoII enrichment were enhanced in regions where Toll-2 was highly enriched, such as the neck and the head tissue near the midline (Figure 3A, B; Figure S4A). These characteristics of *Toll-2^RNAi^* clones agreed with a Toll-2 role in cell sorting and adhesion (Dahmann and Basler, 2000; Fischer et al., 2024). We found that *Toll-2^RNAi^* clones did not trigger the loss of Toll-2:GFP at the interface between control and *Toll-2^RNAi^* cells (Figure 3A). Toll-8 is a known binding partner of Toll-2 (Paré et al., 2014). However the function of Toll-2 in cell sorting and adhesion is unlikely to be mediated by Toll-8 since *Toll-8^RNAi^* clones in the neck or head were not characterized by a smooth boundary nor a loss of Toll-2::GFP at their interface with control cells (Figure S4B) – results in agreement with the lack of Toll-8 function in shaping the *Dfd+* compartment during larval to pupal transition.

**Figure 3:**
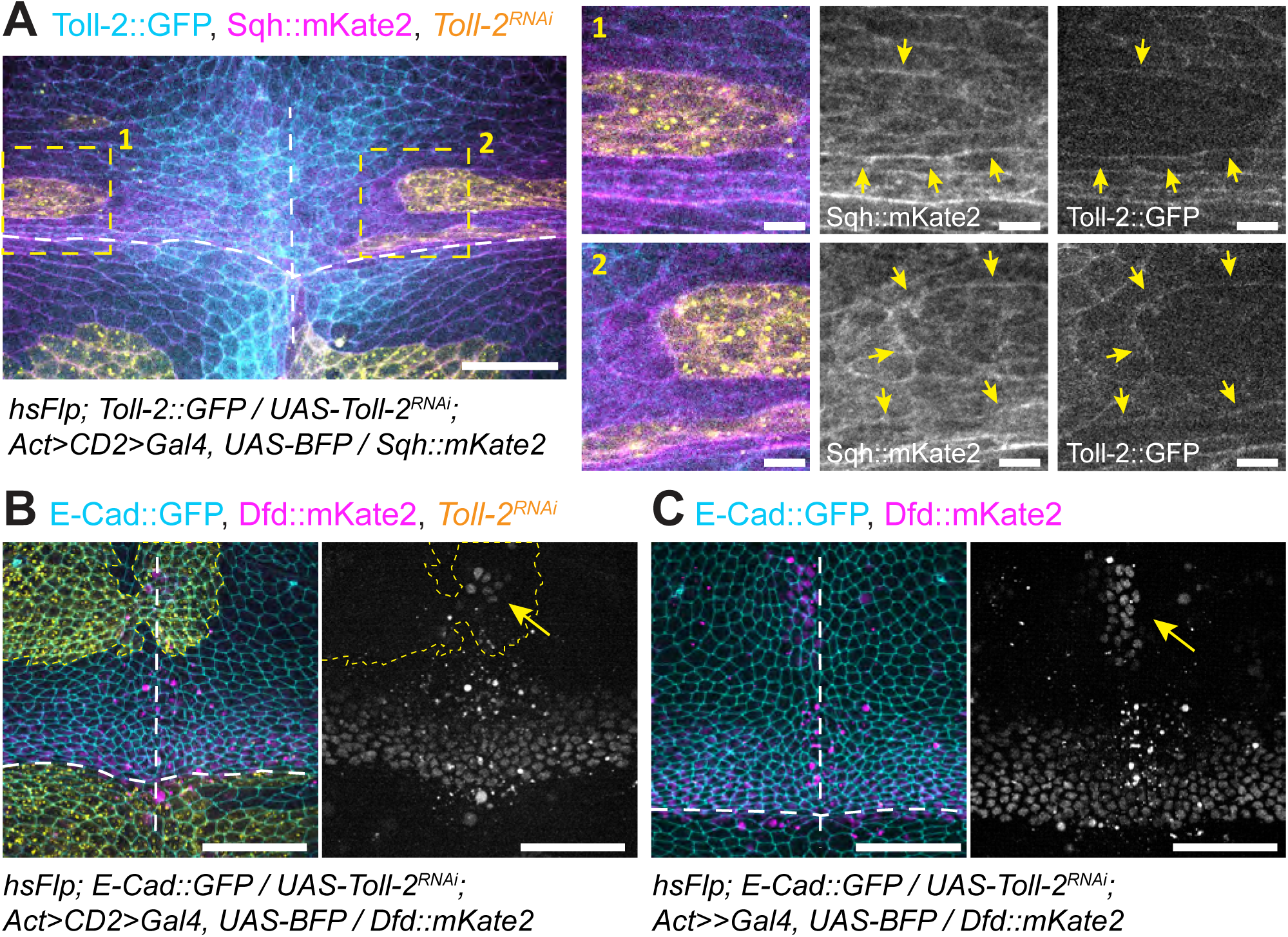
Loss of Toll-2 leads to aberrant segregation of *Dfd-*expressing cells. A: Toll-2::GFP and Sqh::mKate2 in *Toll-2^RNAi^*-expressing clones (at 16 hAPF). White dashed lines mark midline and head/thorax boundary. Yellow dashed boxes mark regions magnified in the right panels. Yellow arrows point at MyoII cable formed around clone (medium panels) or Toll-2 present at the cell junctions of the wildtype cells neighboring the clone (right panels). Scale bar = 50 μm. B: Dfd:mKate2 in neck region with clonal *UAS-Toll-2^RNAi^* expression (at 14h APF). White dashed lines mark midline and head/thorax boundary. Yellow dashed line marks boundaries of the clone with Dfd::mKate2^+^ cells. Arrow points at Dfd::mKate2^+^ cells that sorted into the head tissue. Misplacement of Dfd::mKate2^+^ cells to the head tissue is seen in 5 out of 7 mosaic animals. Scale bars = 50 μm. C: Dfd:mKate2 in neck region with ubiquitous *UAS-Toll-2^RNAi^* expression, driven by tissue-wide flip-out of an *Act>CD2>Gal4* cassette (at 14h APF). White dashed lines mark midline and head/thorax boundary. Arrow points at Dfd::mKate2^+^ cells that sorted into the head tissue. Scale bars = 50 μm.

Having found that loss of Toll-2 function led to defects in the shape of *Dfd+* compartment and the sorting of *Toll-2^RNAi^* cells, we explored whether Toll-2 was necessary to maintain *Dfd+* cells together. Towards this goal, we generated larger *Toll-2^RNAi^* clones in animals where *Dfd+* cells were marked using an in-frame mKate2 knock-in at *Dfd* locus (Dfd::mKate2) (Figure 3B, C). Both in clones encompassing part of the neck or most of the head, neck and thorax epithelia, the clonal loss of Toll-2 function were seen to lead to *Dfd+ Toll-2^KD^* cells sorting away from the neck compartment and into the neighboring medial head tissue (Figure 3B, C). Relevantly, *Toll-2^RNAi^* clones were hard to recover in the medial part of the neck, possibly because they were segregated towards the medial head or lateral neck regions, further underlining the position-dependent response to loss of Toll-2 function. Based on these results, we concluded that Toll-2 is necessary to control the shape of the *Dfd+* homeotic compartment and to maintain *Dfd+* cells together during metamorphosis.

### Medial-to-lateral tissue flows and cell rearrangements characterize pupal *Dfd+* compartment morphogenesis prior to tissue folding onset

The eversion of the head at 12 h APF marks the transition from pre-pupal to pupal stages (Bainbridge and Bownes, 1981). An additional 6-hour time window separates head eversion and the eventual onset of neck folding at 18 h APF, defining a first pupal phase of *Dfd+* compartment development that proceeds its folding. Tracking the evolution of *Dfd>LifeAct::Ruby* neck compartments over this time window revealed that compartment thickness was reduced along the a-p axis while each symmetric half underwent in-plane medial to lateral flows starting at 16 hAPF as measured by particle image velocimetry (PIV) (Figure 4A-C). By segmenting and tracking cells we observed that this early phase of neck morphogenesis was concomitant to a steady increase of cell shape anisotropy along the m-l direction till 18h APF, time at which cell division occurred (Sup Fig 5A-C). Interestingly, the symmetric medial-to-lateral flows were also preceded by a high rate of both cell delaminations and rearrangements, which then accompanied the tissue flow albeit at a lower rate (Figure 4D, E). Notably, the high rate of cell delamination was mainly taking place in the medial region of the *Dfd+* compartment (Figure 4D). We also analyzed the orientation of cell rearrangements and found that till 17 hAPF cell rearrangements mainly produced new m-l oriented junctions consistent with the fact that cell anisotropy increased along the m-l axis. To better understand the contribution of each of these cellular dynamics to the morphogenesis of the *Dfd+* compartment, we then combined cell tracking with a multiscale framework that relates each cell behavior to tissue deformation (Guirao et al., 2015). This analysis revealed that from 15 hAPF onwards, the *Dfd+* neck compartment underwent a convergent-extension deformation leading to its extension along the m-l axis (Figure 4F). Interestingly, this compartment extension was mainly contributed by m-l oriented cell shape elongation. In fact, and in agreement with the higher frequency of cell rearrangements producing m-l junctions, the analysis indicated that cell rearrangements negatively contributed to tissue extension. The analysis also showed that cell delamination has a minimal contribution to tissue m-l extension. Taken together, these results revealed that, prior to its out-of-plane folding, the *Dfd+* compartment underwent an in-plane symmetric medial to lateral tissue flow preceded by both cell rearrangement and delamination. Furthermore, this pre-fold tissue flow is associated with an m-l extension of the compartment, driven mainly by m-l cell elongation, and initially opposed by cell rearrangements.

**Figure 4:**
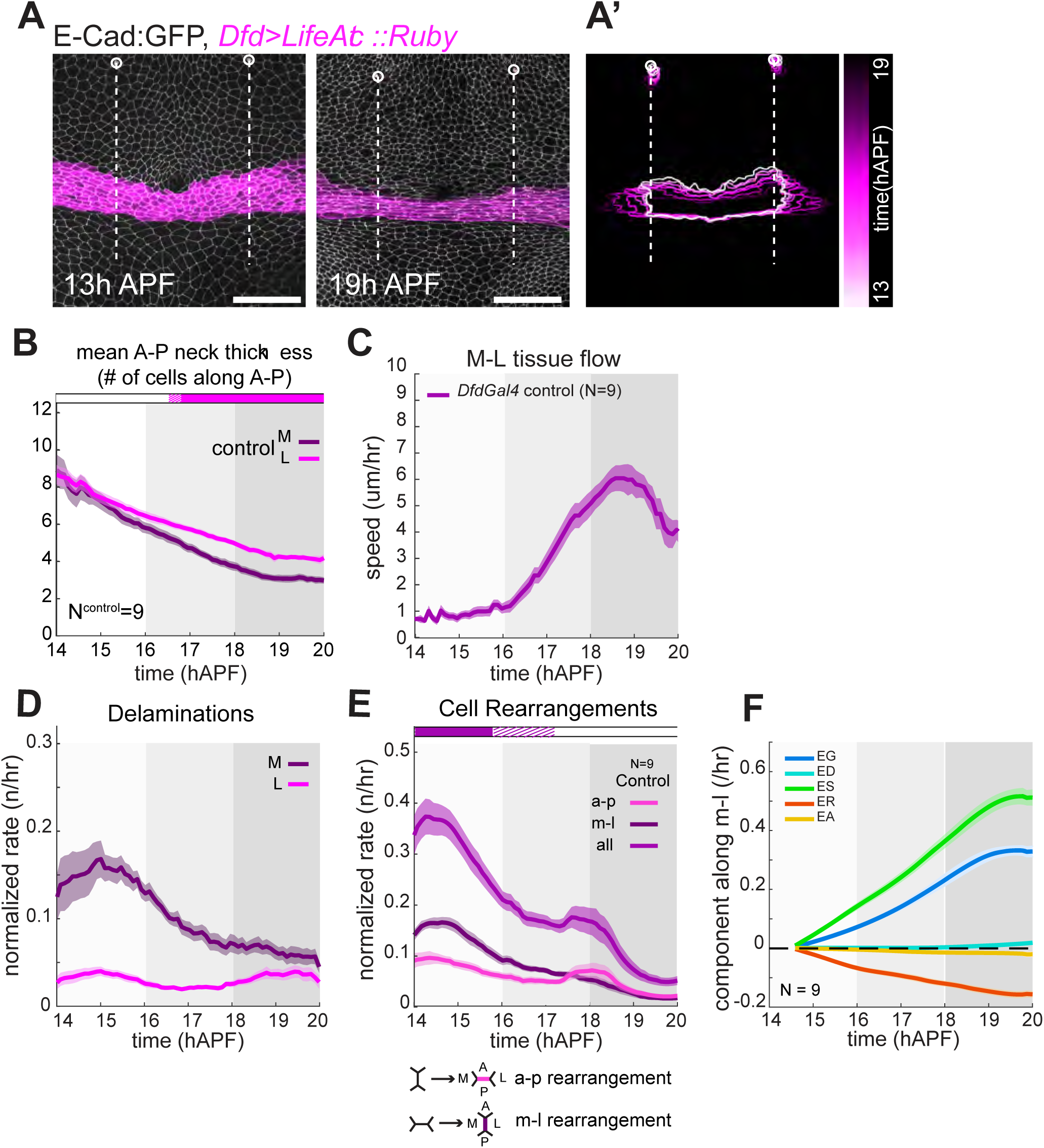
Pupal Dfd compartment morphogenesis prior to fold onset. A: Neck shape as described by *DfdGal4>UAS-LifeAct::Ruby* expression at 13h and 19h APF. White circles mark positions of head macrocaetes used as reference to define medial and lateral neck regions (See Figure 2), white lines correspond to position of macrochaetes along the m-l axis, limiting the initial size of neck masks used in A’. (A’) Progressive shape change of a neck mask defined by *DfdGal4>UAS-LifeAct::Ruby* expressing region between head macrochaetes at 13h APF, segmented and tracked over time. Image shows overlay of neck masks tracked at every hour, from 13h to 19h APF. Scale bars = 50 µm. B: Average number of cells (mean ± sem) found along the anterior-to-posterior (a-p) direction in medial (dark purple) and lateral (magenta) *DfdGal4>UAS-LifeAct::Ruby* expressing Dfd compartments over time. Horizontal boxes: p-values of Welch tests performed between data for medial and lateral regions at successive time points (white p >0.05, striped p<0.05, solid p <0.01). N=9 pupae. C: Plot showing mean rates of medial-to-lateral tissue flow, as measured by PIV, within the *Dfd+* compartmens of *DfdGal4>UAS-LifeAct::Ruby* control animals. N = 9 pupae. D: Mean delamination rates within the medial or lateral *Dfd+* compartment regions of *DfdGal4> UAS-LifeAct::Ruby* control animals, normalized to the total number of cells within analyzed region. N = 9 pupae. E: Mean cell rearrangement rates for all rearrangements, or rearrangements generating junctions in the m-l or a-p orientation within the *Dfd+* compartment of *DfdGal4>UAS-LifeAct::Ruby* control animals, normalized to the total number of cells within analyzed region. Horizontal boxes: p-values of Welch tests performed between data for m-l and a-p rearrangement rates at successive time points (white p >0.05, striped p<0.05, solid p <0.01). N = 9 pupae. F: Plots showing mean tissue constriction/elongation (EG) and the average contribution of cell divisions (ED), shape changes (ES), rearrangements (ER), and delaminations (EA) to this total morphogenetic change, over time, for the Dfd compartments of *DfdGal4>UAS-LifeAct::Ruby* control animals (mean ± sem). N = 9 pupae. For all plots, period between onset of medial-to-lateral tissue flow and onset of folding (16h to 18h APF) shown in medium grey background; period after onset of invagination (18h APF onwards) shown in dark grey background. Reference time points are based on observed average tissue movement initiation times in controls. All quantities plotted are moving averages over a 2h sliding window, unless otherwise stated.

### Toll-2 contributes to neck morphogenesis via regulating cell delaminations and rearrangements

We then analyzed whether and how Toll-2 contributes to neck flow and morphogenesis by comparing control and *Dfd>Toll-2^RNAi^* neck dynamics. We found that the medial to lateral tissue flow started significantly earlier in *Dfd>Toll-2^RNAi^* animals (Figure 5A). Interestingly, the earlier in-plane tissue flow was also associated with an earlier onset of neck folding as characterized by the earlier deepening of the tissue without changes in the rate of tissue deepening speed (Figure 5A-C). To best illustrate these changes in neck dynamics, we plotted the neck depth position in *Dfd>Toll-2^RNAi^* animal versus control animal for each time-point. Fitting this set of points by a line showed that its intercept with y-axis is positive consistent with an advanced timing of deepening while its slope is closed to 1 in agreement with a similar deepening speed (Figure 5D). Therefore, the neck starts to fold earlier, but not faster in *Dfd>Toll-2^RNAi^* animals. We have previously shown that the speed of folding is determined by the tissue tension estimated by laser ablation (Villedieu et al., 2023) (Figure S5D-K). In agreement with the lack of change in folding speed, neither Myo II levels nor m-l tension estimated by laser ablation were altered in *Dfd>Toll-2^RNAi^* animals (Figure 3A and Figure S5L). Therefore, without affecting tissue tension, the *Dfd+* compartment initiated an earlier in-plane flow followed by an earlier deepening of the compartment in *Dfd>Toll-2^RNAi^* animals. Importantly, we found that the earlier onset of neck folding was accompanied by a restoration of the medial neck thickness in *Dfd>Toll-2^RNAi^* animals (Figure 5E-G).

**Figure 5:**
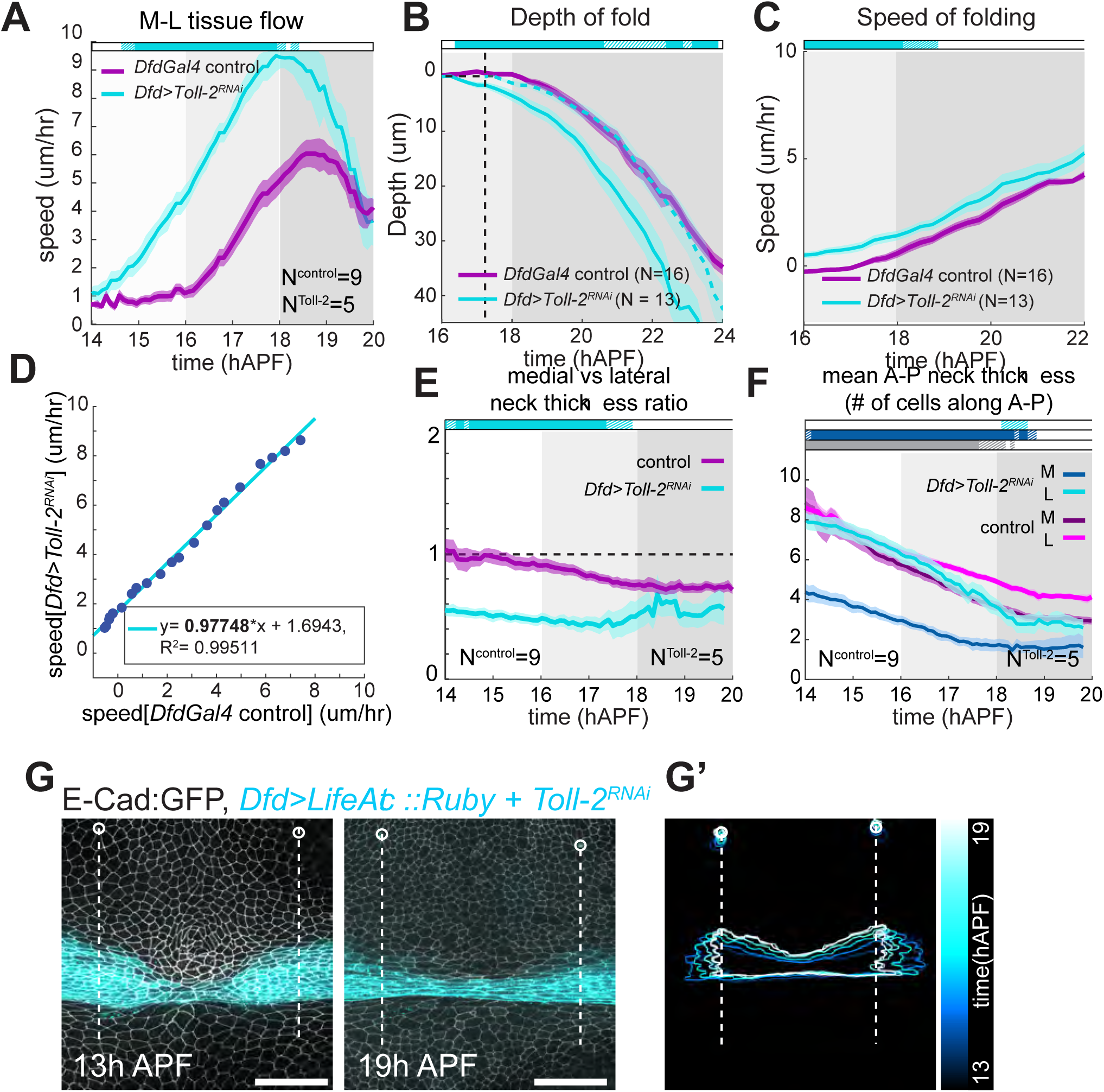
Loss of Toll-2 function impacts timing of tissue movements. A: Plot showing mean rates of tissue flow along the medial-lateral axis, as measured by PIV, within the *Dfd+* neck region of *DfdGal4>UAS-Toll-2^RNAi^* (cyan) and *DfdGal4>UAS-LifeAct::Ruby* control (purple) animals. Horizontal box: p-values of Welch tests performed between control and experimental conditions at successive time points (white p >0.05, striped p<0.05, solid p <0.01). *DfdGal4>UAS-LifeAct::Ruby* control N = 9 pupae; *DfdGal4>UAS-Toll-2^RNAi^* N = 5 pupae. B: Plot showing mean depth of neck fold over time, in *DfdGal4>UAS-Toll-2^RNAi^* (cyan) and *DfdGal4>UAS-LifeAct::Ruby* control (purple) animals. Dashed cyan line marks the fold depth curve for *DfdGal4>UAS-Toll-2^RNAi^*, shifted in time. Notice that while the inflection point for the two curves are different along the time axis, the overall shape of the curve is quite similar (see also, Fig Sup 5). Horizontal box: p-values of Welch tests performed between control and experimental conditions at successive time points (white p >0.05, striped p<0.05, solid p <0.01). *DfdGal4>UAS-LifeAct::Ruby* control N = 18 pupae; *DfdGal4>UAS-Toll-2^RNAi^* N = 14 pupae. C: Plot showing mean speed of fold deepening over time, in *DfdGal4>UAS-Toll-2^RNAi^* (cyan) and *DfdGal4>UAS-LifeAct::Ruby* control (purple) animals. Horizontal box: p-values of Welch tests performed between control and experimental conditions at successive time points (white p >0.05, striped p<0.05, solid p <0.01). *DfdGal4>UAS-LifeAct::Ruby* control N = 18 pupae; *DfdGal4>UAS-Toll-2^RNAi^* N = 14 pupae. D: Mean speed of fold deepening in *DfdGal4>UAS-Toll-2^RNAi^* plotted against mean speed of fold deepening in *DfdGal4>UAS-LifeAct::Ruby* control for each corresponding time point. *DfdGal4>UAS-LifeAct::Ruby* control N = 18 pupae; *DfdGal4>UAS-Toll-2^RNAi^* N = 14 pupae. E: Average ratio of medial to lateral a-p neck thicknesses (mean ± sem) over time, in *DfdGal4>UAS-LifeActRuby* control and *DfdGal4> LifeActRuby + UAS-Toll-2^RNAi^* expressing animals (corresponding to Figures 4B and 5G, respectively). Horizontal box: p-values of Welch tests performed between control and experimental conditions at successive time points (white p >0.05, striped p<0.05, solid p <0.01). *DfdGal4>UAS-LifeAct::Ruby* control, N=9 pupae; *DfdGal4>UAS-Toll-2^RNAi^*, N=5 pupae. F: Average number of cells (mean ± sem) found along the a-p direction in medial (dark shades) and lateral (light shades) *DfdGal4>UAS-LifeAct::Ruby* expressing Dfd compartments over time in control (purple) and *DfdGal4>UAS-Toll-2^RNAi^* (blue) animals. Horizontal boxes: p-values of Welch tests performed between data for control and *Toll-2^RNAi^* medial (cyan) or lateral (dark blue) regions, or comparing medial and lateral regions of *Toll-2^RNAi^* necks, at successive time points (white p >0.05, striped p<0.05, solid p <0.01). *DfdGal4>UAS-LifeAct::Ruby* control, N=9 pupae; *DfdGal4>UAS-Toll-2^RNAi^*, N=5 pupae. G: Neck shape as described by *DfdGal4>UAS-LifeAct::Ruby* expression, in a *DfdGal4>UAS-Toll-2^RNAi^* animal at 13h and 19h APF. White circles mark positions of head macrocaetes used as reference to define medial and lateral neck regions (See Figure 2), white lines correspond to position of macrochaetes along the m-l axis, limiting the initial size of neck masks used in G’. (G’) Progressive shape change of a neck mask defined by *DfdGal4>UAS-LifeAct::Ruby* expressing region between head macrochaetes in a *DfdGal4>UAS-Toll-2^RNAi^* animal at 13h APF, segmented and tracked over time. Image shows overlay of neck masks tracked at every hour, from 13h to 19h APF. Scale bars = 50 µm. For all plots, period between onset of medial-to-lateral tissue flow and onset of folding (16h to 18h APF) shown in medium grey background; period after onset of invagination (18h APF onwards) shown in dark grey background. Reference time points are based on observed average tissue movement initiation times in controls. All quantities plotted are moving averages over a 2h sliding window, unless otherwise stated.

Several TLRs, including Toll-2 are shown to control cell rearrangements (Paré et al., 2014; Tamada et al., 2021), and cell death in developing tissues (Fischer et al., 2024; Meyer et al., 2014). To understand how Toll-2 promotes the earlier onset of morphogenesis we compared cell dynamics in control and *Dfd>Toll-2^RNAi^* animals. We found that *Dfd>Toll-2^RNAi^* animals were characterized by a lower delamination rate in the medial region of the neck compartment, while cell division rate was largely unaffected (Figure 6A and Figure S5C). In agreement with a putative role in controlling cell delamination, Toll-2::GFP is enriched in cells that eventually underwent delamination (Figure 6B; Movie 4). In addition, in *Dfd>Toll-2^RNAi^* animals the rate of cell rearrangements, specifically those leading to the formation of m-l junctions, was strongly reduced (Figure 6C). Accordingly, we could observe Toll-2 to be relatively more enriched at cell junctions forming upon cell rearrangements (Figure 6D). We then compared the morphogenesis of the *Dfd+* compartment in control and *Dfd>Toll-2^RNAi^* animals (Figure 6E). This revealed that *Dfd>Toll-2^RNAi^* were characterized by a stronger m-l extension of the tissue associated with the decreased rate of cell rearrangements opposing m-l tissue extension. Altogether these results supported a role for Toll-2 in controlling the timing of tissue flow along the m-l axis and the morphogenesis of the neck compartment. These defects in global tissue flow and morphogenesis were associated with a function of Toll-2 in promoting cell delamination and m-l cell rearrangements that oppose tissue extension.

**Figure 6:**
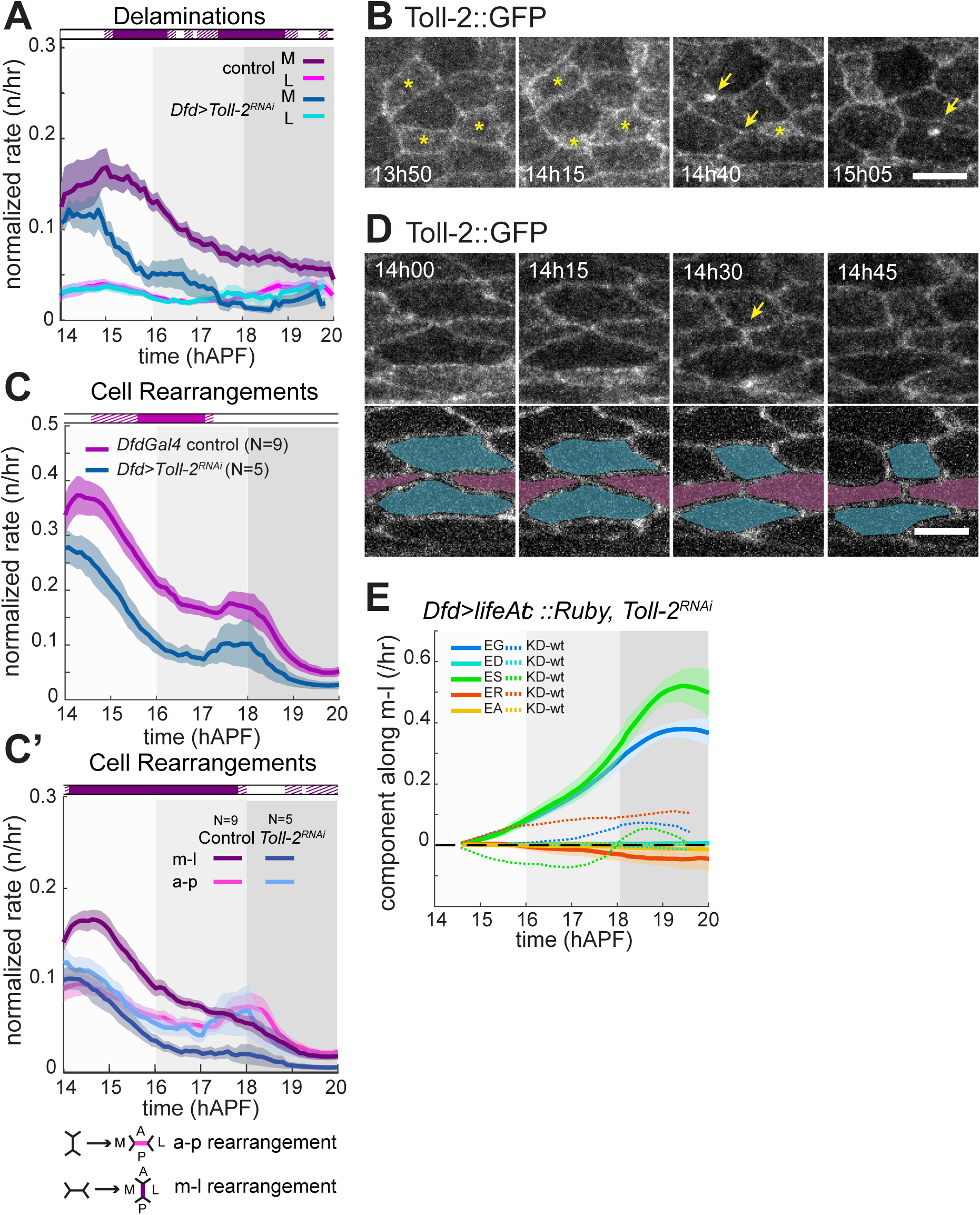
Toll-2 regulates cell delamination and rearrangements in the early pupal phase. A: Mean delamination rates within the medial (darker shades) or lateral (lighter shades) *Dfd+* compartment regions of *DfdGal4> UAS-LifeAct::Ruby* control (purple) and *DfdGal4> UAS-LifeAct::Ruby + UAS-Toll-2^RNAi^* (blue) animals, normalized to the total number of cells within analyzed region. Horizontal box: p-values of Welch tests performed between control and experimental medial regions at successive time points (white p >0.05, striped p<0.05, solid p <0.01). *DfdGal4>UAS-LifeAct::Ruby* control, N=9 pupae; *DfdGal4>UAS-Toll-2^RNAi^*, N=5 pupae. B: Close ups of time-lapse images of Toll-2::GFP in cells that eventually delaminate, prior to and during delaminations (images taken from Movie 4). Arrows point at positions of newly delaminated cells. Yellow asterisks mark cells going through delamination. Notice how Toll-2::GFP levels in these cells are higher than their neighbors, even before their apical cell area is reduced. Scale bar = 10 μm. C: Mean cell rearrangement rates within the *Dfd+* compartment of *DfdGal4>UAS-LifeAct::Ruby* control (magenta) and *DfdGal4>UAS-LifeAct::Ruby + UAS-Toll-2^RNAi^* (cyan) animals, normalized to the total number of cells within analyzed region. Horizontal boxes: p-values of Welch tests performed between control and experimental conditions at successive time points (white p >0.05, striped p<0.05, solid p <0.01). (C’) m-l (dark shades) or a-p (light shades) cell rearrangement rates within the *Dfd+* compartment of *DfdGal4>UAS-LifeAct::Ruby* control (purple) and and *DfdGal4> UAS-LifeAct::Ruby + UAS-Toll-2^RNAi^* (blue) animals, normalized to the total number of cells within analyzed region. m-l or a-p rearrangements refer to rearrangements that form a new junction along the m-l or a-p axis, respectively. Horizontal boxes: p-values of Welch tests performed between data for m-l rearrangement rates in control and *Toll-2^RNAi^* conditions at successive time points (white p >0.05, striped p<0.05, solid p <0.01). *DfdGal4>UAS-LifeAct::Ruby* control, N=9 pupae; *DfdGal4>UAS-Toll-2^RNAi^*, N=5 pupae. D: Time-lapse images showing Toll-2::GFP enrichment in a cell junction undergoing an m-l rearrangement (images taken from movie 4). Arrow points at junction with Toll-2 enrichment. Bottom panels show same images with the cells undergoing rearrangement highlighted for easy visualization of the transition. Scale bar = 10 μm. E: Plots showing mean tissue constriction/elongation (EG) and the average contribution of cell divisions (ED), shape changes (ES), rearrangements (ER), and delaminations (EA) to this total morphogenetic change, over time, for the Dfd compartments of *DfdGal4>UAS-LifeAct::Ruby + UAS-Toll-2^RNAi^* animals (mean ± sem). Dashed lines show difference from *DfdGal4>UAS-LifeAct::Ruby* control values shown in Figure 4F. *DfdGal4>UAS-LifeAct::Ruby* control, N=9 pupae; *DfdGal4>UAS-Toll-2^RNAi^*, N=5 pupae. For all plots, period between onset of medial-to-lateral tissue flow and onset of folding (16h to 18h APF) shown in medium grey background; period after onset of invagination (18h APF onwards) shown in dark grey background. Reference time points are based on observed average tissue movement initiation times in controls. All quantities plotted are moving averages over a 2h sliding window, unless otherwise stated.

### Apoptosis and cell rearrangement control the initial tissue flow and the onset of folding

The timing of cell delamination preceded and coincided with both high rates of cell rearrangements and the m-l tissue flow. We therefore explored their contribution to *Dfd+* compartment development by expressing the apoptosis inhibitor *Diap1* within the compartment (*Dfd>Diap1*) and fully characterizing the tissue dynamics by PIV as well as cell segmentation and tracking. This first revealed a drastic decrease in cell delamination indicating that the observed delamination in the neck corresponded to cell apoptosis (Figure 7A). Characterizing the morphogenesis of *Dfd>Diap1* necks we found that, as observed in *Dfd>Toll-2^RNAi^* animals, the medial to lateral compartment flow occurs earlier and at a faster speed – though less pronounced than that seen in *Dfd>Toll-2^KD^* animals (Figure 7B). Interestingly the reduced rate of apoptosis was also associated with an initial decrease of the rate of rearrangements leading to the formation of m-l junctions (Figure 7C). Furthermore, in agreement with the fact that apoptosis did not contribute to the convergence-extension of the tissue in controls we observed that the morphogenesis of the tissue was similar in *Dfd>Diap1* and control animal (Figure 7D). To test a putative role of cell rearrangement in the regulation of the early m-l flow of the tissue, we reduced the function of Xit, an endoplasmic reticulum glucosyltransferase known to promote junction elongation during cell rearrangements in the early *Drosophila* embryo (Zhang et al., 2014). We found that *Dfd>Xit^RNAi^* animals were characterized by an earlier onset of neck deepening (Figure 7E, Figure S6 A,B). Together this suggested that Toll-2 acts in part by promoting cell apoptosis and m-l rearrangement to prevent earlier onset of tissue flow and neck folding.

**Figure 7:**
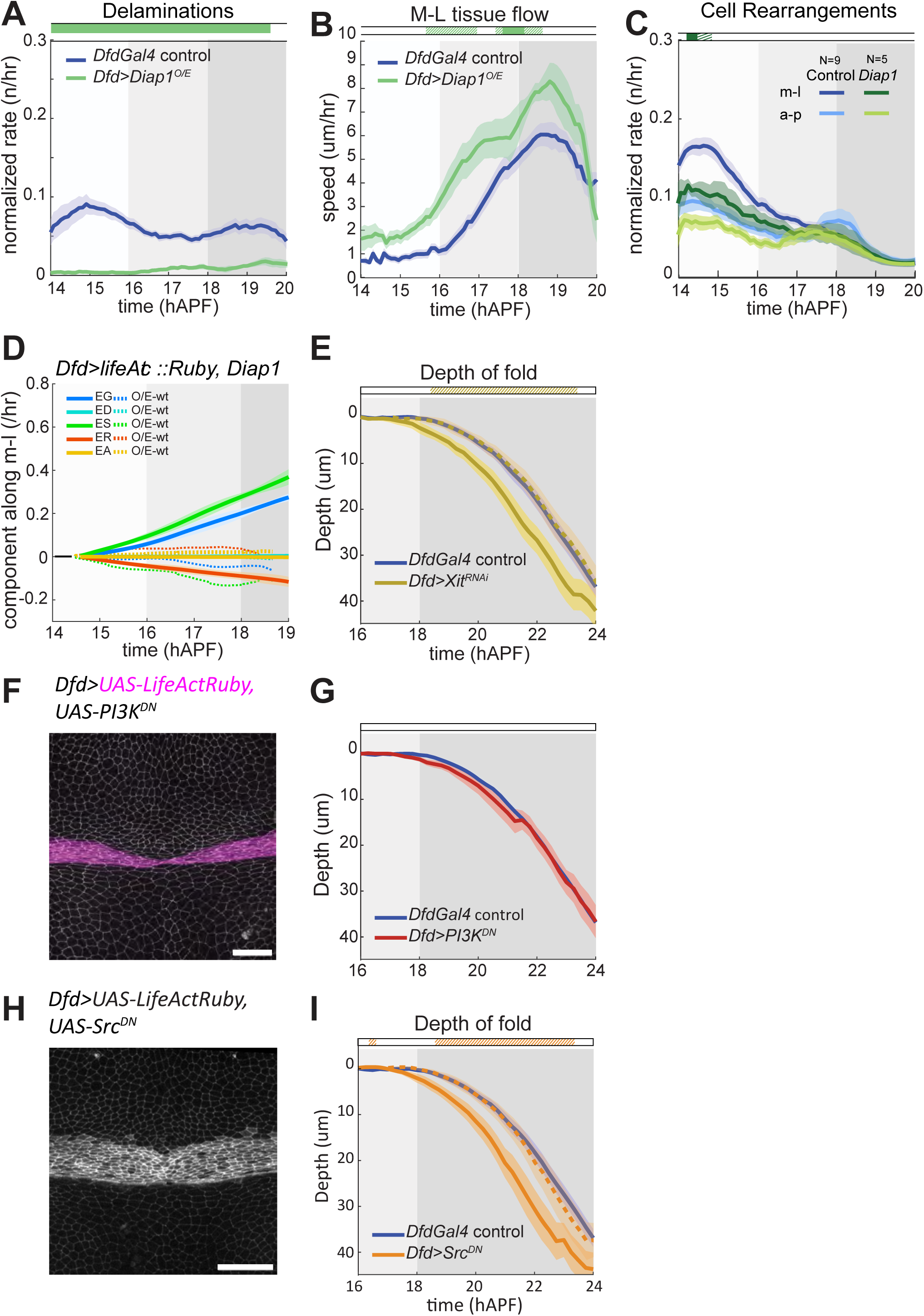
Reducing delamination and rearrangement rates alter tissue flow and fold dynamics. A: Mean total delamination rates within the *Dfd+* compartment regions of *DfdGal4> UAS-LifeAct::Ruby* control (blue) and *DfdGal4> UAS-LifeAct::Ruby + UAS-Diap1* (green) animals, normalized to the total number of cells within analyzed region. Horizontal box: p-values of Welch tests performed between control and experimental medial regions at successive time points (white p >0.05, striped p<0.05, solid p <0.01). *DfdGal4>UAS-LifeAct::Ruby* control N = 9; *DfdGal4> UAS-LifeAct::Ruby + UAS-Diap1* N = 4 pupae. B: Plot showing mean rates of tissue flow along the medial-lateral axis, as measured by PIV, within the Dfd compartment of *DfdGal4>UAS-LifeAct::Ruby* control (blue) and *DfdGal4> UAS-LifeAct::Ruby* + *UAS-Diap1* (green) animals. Horizontal box: p-values of Welch tests performed between control and experimental conditions at successive time points (white p >0.05, striped p<0.05, solid p <0.01). *DfdGal4>UAS-LifeAct::Ruby* control N = 9 pupae; *DfdGal4> UAS-LifeAct::Ruby + UAS-Diap1* N = 4 pupae. C: Mean m-l (dark shades) or a-p (light shades) cell rearrangement rates within the *Dfd+* compartment of *DfdGal4>UAS-LifeAct::Ruby* control (blue) and *DfdGal4>UAS-LifeAct::Ruby + UAS-Diap1* (green) animals, normalized to the total number of cells within analyzed region. m-l or a-p rearrangements refer to rearrangements that form a new junction along the m-l or a-p axis, respectively. Horizontal boxes: p-values of Welch tests performed between data for m-l rearrangement rates in control and experimental conditions at successive time points (white p >0.05, striped p<0.05, solid p <0.01). *DfdGal4>UAS-LifeAct::Ruby* control N = 9; *DfdGal4> UAS-LifeAct::Ruby + UAS-Diap1* N = 4 pupae. D: Plots showing mean tissue constriction/elongation (EG) and the average contribution of cell divisions (ED), shape changes (ES), rearrangements (ER), and delaminations (EA) to this total morphogenetic change, over time, for the Dfd compartments of *DfdGal4>UAS-LifeAct::Ruby + UAS-Diap1* animals (mean ± sem). Dashed lines show difference from *DfdGal4>UAS-LifeAct::Ruby* control values shown in Figure 4F. *DfdGal4>UAS-LifeAct::Ruby* control N = 9; *DfdGal4> UAS-LifeAct::Ruby + UAS-Diap1* N = 4 pupae. E: Plot showing mean depth of neck fold over time, in *DfdGal4>UAS-Xit^RNAi^* (yellow) and *DfdGal4* control (blue) animals. Dashed yellow line marks the fold depth curve for *DfdGal4> UAS-Xit^RNAi^*, shifted in time. Horizontal box: p-values of Welch tests performed between control and experimental conditions at successive time points (white p >0.05, striped p<0.05, solid p <0.01). *DfdGal4* control N = 18 pupae; *DfdGal4> UAS-Xit^RNAi^* N = 12 pupae. F: Representative image showing *Dfd+* compartment shape in a *DfdGal4>UAS-LifeAct::Ruby + UAS-PI3K^DN^* animal at 14h APF. Scale bar = 50 μm. G: Plot showing mean depth of neck fold over time, in *DfdGal4>UAS-LifeAct::Ruby + UAS-PI3K^DN^* (red) and . *DfdGal4>UAS-LifeAct::Ruby* control (blue) animals. Horizontal box: p-values of Welch tests performed between control and experimental conditions at successive time points (white p >0.05, striped p<0.05, solid p <0.01). *DfdGal4>UAS-LifeAct::Ruby* control N = 16 pupae; *DfdGal4>UAS-PI3K^DN^* N = 10 pupae. H: Representative image showing *Dfd+* compartment shape in a *DfdGal4>UAS-Src^DN^::Venus* animal at 14h APF. Scale bar = 50 μm. I: Plot showing mean depth of neck fold over time, in *DfdGal4>UAS-Src^DN^* (orange) and *DfdGal4* control (blue) animals. Dashed orange line marks the fold depth curve for *DfdGal4>UAS-Src^DN^*, shifted in time. Horizontal box: p-values of Welch tests performed between control and experimental conditions at successive time points (white p >0.05, striped p<0.05, solid p <0.01). *DfdGal4* control N = 18 pupae; *DfdGal4>*

### Toll-2 acts via distinct effectors to regulate the initial shape and dynamics of the Dfd homeotic compartment

So far, our results identified that Toll-2 controls both the initial shape of the *Dfd+* compartment and the temporality of neck folding. To determine whether the initial defects in the shape of the *Dfd+* compartment promote the earlier onset of neck morphogenesis we decided to analyze the contribution of known effectors Toll-2, PIK3 and Src that have been reported to act downstream of Toll-2 during *Drosophila* germband elongation (Tamada et al., 2021). To abrogate PIK3 function, we over-expressed a dominant negative form of PI3K in the *Dfd+* compartment (*Dfd>PI3K^DN^*). This resulted in a defect in neck compartment shape similar to the one observed in of *Dfd>Toll-2^RNA^* since *Dfd>PI3K^DN^* necks are narrower in the medial part (Figure 7F). Strikingly we found that in *Dfd>PI3K^DN^* animals neck folding occurred similarly to control animals (Figure 7G). In contrast, *Dfd>Src^DN^* necks were not defected in shape, while they exhibited an early folding phenotype (Figure 7H, I; Figure S6 C, D). Interestingly *Dfd>Src^DN^* tissue were also characterized by an increased deepening speed during folding (Figure 7I; Figure S6 C, D) as observed upon increased MyoII activity (Figure S5 F, I) (Villedieu et al., 2023). Together, these results suggest that the initial shape and later fold timing phenotypes can be uncoupled, and that the latter is unlikely to be merely a result of the first.

Collectively our results show a dual role for Toll-2 in the morphogenesis of the *Dfd* homeotic domain via two distinct effectors PI3K and Src. Early on, Toll-2 is required at the time of disc fusion, to correctly sort the left and right populations of *Dfd+* neck cells into a uniform tissue compartment. Following head eversion, Toll-2 regulates cell apoptosis and cell rearrangements in an early phase of pupal neck morphogenesis to regulate the onset of neck m-l flow and folding.

## Discussion

Tissue compartmentalization by Hox genes is an evolutionarily conserved feature of animal development and is crucial for the patterning, growth, and shaping of the animal body (Hubert and Wellik, 2024). Here, we explored the mechanisms of homeotic compartment formation and morphogenesis, focusing on the contribution of Toll-like receptors (TLRs) to these processes. By characterizing the formation of the pupal *Dfd+* compartment and its cellular dynamics from the onset of morphogenesis, we identified the essential role of Toll-2 in the proper shaping of the pupal *Dfd+* compartment and the sorting of *Dfd+* cells. Furthermore, our detailed quantitative analysis of pupal *Dfd+* compartment morphogenesis uncovered an unexpected role for Toll-2 in timing tissue flows and folding by regulating apoptosis and cell rearrangement rates. Thus, our findings provide novel insights into homeotic compartment development and its spatiotemporal regulation by TLRs.

As in all holometabolous insects, the *Drosophila* adult is shaped by the fusion of larval imaginal tissues during metamorphosis. Numerous studies have highlighted the importance of signaling pathways, such as JNK, Dpp, and Fat/Dachsous, and the contributions of the larval epidermal cell layer in fusing the mesothoracic wing discs along the dorsal midline (Athilingam et al., 2022; Lu et al., 2022; Martín-Blanco et al., 2000; Usui and Simpson, 2000; Zeitlinger and Bohmann, 1999). However, the internal positioning of the eye-antennal discs, enclosed by the thorax prior to head eversion, has rendered their pre-pupal morphogenesis elusive. Using *in vitro* cultured eye-antennal discs, the fusion of these discs has been described as achieving perfect alignment and symmetry of patterned head structures along the line of fusion (Milner et al., 1984). By characterizing the *in vivo* development of the *Dfd+* compartment during eye-antennal disc fusion, we delineated the complex 3D morphogenetic choreography that aligns matching bilateral tissue regions to define the a-p compartment pattern of the pupa. These *in vivo* findings extend previous indirect characterizations of the origin of adult head structures. Indeed, the peripodial epithelium of the eye-antennal imaginal disc was suggested to contribute to adult head structures based on fate maps from transplantation experiments (Haynie and Bryant, 1986) and adult expression of a peripodia-specific transcriptional reporter (Bessa and Casares, 2005; Lee et al., 2007). Accordingly, *Dfd* was previously reported to be expressed in eye-antennal peripodial cells (Diederich et al., 1991), with this expression shown to be important for signaling to the disc proper (Stultz et al., 2012). By tracking *Dfd+* cells from their peripodial origin to the presumptive adult epithelium, we opened novel research avenues into the mechanisms regulating the peripodia-to-primordium transition and head structure formation in holometabolous insects.

The misshapen pupal neck phenotype observed in the absence of Toll-2 points to specific mechanisms for fusing, aligning, and matching tissue compartment patterns. Precise compartmental alignment about the midline has been reported for thoracic and abdominal segments and was proposed to occur via cell-cell recognition of positional values (Usui and Simpson, 2000). Our analysis of Toll-2 function and previous findings suggest that Toll-2 may act through diverse mechanisms. TLRs can regulate actomyosin dynamics to control cell behaviors during morphogenesis, wound healing, or tissue homeostasis (Carvalho et al., 2014; Fischer et al., 2024; Lavalou et al., 2021; Paré et al., 2014; Tamada et al., 2021; Villedieu et al., 2023). It is conceivable that Toll-2 promotes cell movement to bring left and right *Dfd+* compartments together via actomyosin regulation. In addition, the contribution of Toll-2 to *Dfd+* compartment formation may relate to the diverse roles of TLRs in cell-cell signaling. For instance, the canonical Toll pathway can interact with JNK signaling (Kawai et al., 2024), a pathway critical for thoracic disc fusion, and Toll-1, -2, and -8 can induce JNK activity in the wing disc (Fischer et al., 2024; Li et al., 2020). The similarity of defects observed in the *Dfd+* compartment upon knockdown of *Cadherin* and *Toll-2* suggests that Toll-2 may directly regulate adhesion during the fusion of bilateral eye-antennal discs. Additionally, the misplacement of *Toll-2^RNAi^*-expressing *Dfd+* cells into the head tissue indicates that Toll-2 not only adheres left and right compartments but also maintains *Dfd+* cell segregation from neighboring head tissue. As Toll-2 is also expressed in the head, it may act in conjunction with other surface receptors to define the positional value of *Dfd+* cells. Alternatively, Toll-2 levels or distinct downstream signaling within the neck region may specify *Dfd+* cell identity. These hypotheses align with previous loss-of-function phenotypes of TLRs, including Toll-2, which are position-dependent (Fischer et al., 2024) . Our findings that *Toll-2^RNAi^* clones separate from their neighbors with a smooth, MyoII-enriched interface suggest that actomyosin regulation contributes to the positional information defined by Toll-2. Exploring Toll-2 function and analyzing PI3K’s role within the *Dfd+* compartment will be instrumental in elucidating potential mechanisms that inform cells of their position and contribute to compartment formation.

In addition to characterizing the mechanisms of *Dfd+* homeotic compartment formation, our exploration of Toll-2 function enhances the understanding of folding regulation. The mechanisms contributing to tissue folding has been extensively explored (Martin and Goldstein, 2014). Yet our understanding of the control of folding timing is far less advanced. Previous studies identified that correct cell size and architecture within a folding domain (Xie et al., 2016) or the completion of blastocyst cellularization (Horo et al., 2024) are necessary to initiate folding. Here, we uncovered a mechanism that delays fold deepening without affecting its speed. Our detailed quantitative study of tissue dynamics in the pupal *Dfd+* compartment indicates that an early phase of frequent cell delamination and rearrangement instructs the timing of subsequent tissue flows and folding. Specifically, cell rearrangements forming medial-lateral (m-l) oriented junctions in this early phase counteract medial-to-lateral tissue flows associated with m-l cell elongation. These rearrangements appear partially dependent on Toll-2 function in inducing apoptosis. Implicit in this model is a link between in-plane flow and out-of-plane folding movements of the *Dfd*+ compartment. It is conceivable that 2D flow movements on the curved neck surface translate into inward motion in 3D. By nature of tissue curvature, lateral regions where flows displace cells are positioned deeper relative to the dorsal midline. Furthermore, lateral extension of the curved tissue surface may increase curvature, further promoting folding (Villedieu et al., 2023).

Toll-2’s role in cell rearrangements may differ between tissues and rely on distinct effectors. During germband extension in the *Drosophila* embryo, planar-polarized Toll-2 localization orients cell rearrangements by polarizing Src and PI3K activity to promote junction contraction, favoring rearrangements that reduce cell anisotropy and thus promoting tissue flows associated with germband elongation (Paré et al., 2014; Tamada et al., 2021) . Here, we found that, in the *Dfd+* compartment, Toll-2 instead promotes cell rearrangements that increase cell shape anisotropy, preventing m-l tissue flow. Such rearrangements promoting cell elongation have been described in the *Drosophila* pupal wing and their regulation in part deciphered (Bardet et al., 2013; Etournay et al., 2015; Ikawa et al., 2023). It will be interesting to explore whether distinct regulations of PI3K and Src explain Toll-2’s varying roles in orienting cell rearrangements and tissue flow during morphogenesis.

TLRs, including Toll-2, mediate cell competition in *Drosophila*, promoting apoptosis to ensure tissue fitness (Alpar et al., 2018; Meyer et al., 2014). While studies of cell competition typically involve ectopically induced genetic heterogeneities, we identified a physiologically occurring phase of Toll-2-dependent apoptosis. Interestingly, the effectors mediating apoptosis may overlap, as Src promotes basal cell delamination during cell competition in MDCK type I cells (Kajiwara et al., 2022). By exploring Toll-2 function, our results suggest a role for apoptosis in promoting rearrangements thereby timing tissue flows. Whether and how apoptosis can orient cell rearrangements remains an intriguing question for future research. Additionally, previous studies have shown the role of cell apoptosis in controlling folding (Monier et al., 2015; Roellig et al., 2022; Villedieu et al., 2023). Together, exploring Toll-2’s role in apoptosis will illuminate the mechanisms that ensure tissue fitness and dynamic regulation during development and homeostasis.

## Acknowledgments

We thank J. Zallen, the Bloomington Drosophila Stock Center, Transgenic RNAi Project at Harvard Medical School, Vienna Drosophila Resource Center, and the Developmental studies Hybridoma Bank for reagents; the department of genetics and developmental biology imaging facility PICT-IBiSA@BDD for help with microscopy; This work was supported by Institut Curie, CNRS, INSERM, ANR (TiMecaDiv 20CE13000801), ANR (ChronoDamage 20CE13-0013), CANCERO-INCA (PLBIO2020/BELLAICHE), ANR Labex DEEP (11-LBX-0044, ANR-10-IDEX-0001-02) and ARC (PDF20181208399) fellowship.

## Author contributions

L.A. and Y.B. designed the project. S.P. produced reagents. L.A. developed experimental methods and performed experiments. R.P., A.L. and M.E. developed data analysis methods. L.A. analyzed the data. L.A and Y.B. wrote the manuscript.

## Competing interests

The authors declare no competing interests.

## Materials and Methods

### Fly husbandry and stocks

A list of *Drosophila melanogaster* stocks used and associated references is provided in Supplementary Table 1. For Dfd compartment-specific or clonal expression of loss-of-function or gain-of-function constructs or fluorescently tagged proteins, the Gal4/UAS system was used in combination with temperature sensitive Gal80^ts^. To control the transcriptional activity of the *Dfd-GAL4* drivers, animals were raised at 18°C and incubated at 29°C for 72h prior to imaging for *Dfd>Toll-2^RNAi^* and *Dfd>Xit^RNAi^*, for 24h for *Dfd>PI3K^DN^* and *Dfd>Src42-DN-Venus (Dfd>Src^DN^)*, or for 8h for *Dfd>Diap1* and *Dfd>shg^RNAi^* conditions. For the screening of TLR and LRR surface molecules, *UAS-RNAi* was expressed in the absence of Gal80^ts^, and flies were raised at 25°C at all times. For clonal loss-of-function analyses the FLP/FRT system was used, larvae were heat-shocked at 37°C for 15min, 72h prior to imaging, and kept at 29°C at all times.

### Generation of Toll-2::GFP

### Mounting and imaging of pupa

#### Time-lapse live imaging

For live imaging of pre-pupal stages, pupae were collected at 0h APF and adhered ventrally on a slide using double-sided tape (Scotch 3M). For better viewing of eye-antennal discs, the anterior ends of the pupae were raised by placing 6 layers of double-sided tape below them. A 0.5 μm drop of water-soluble 2,2’-Thiodiethanol (TDE) mounting medium (Sigma Aldrich) was positioned on the opercular ridge, and a 24×40mm #1 (0.13-0.16mm) coverslip (Knittel Glass) was placed over the pupae, supported by two spacer stacks of six and five 18×18mm #1 (0.13-0.16mm) coverslips (Thermo Scientific, Menzel-Glaser), adhered close to the edges of the slide, posteriorly and anteriorly to the pupae respectively. Care was taken to prevent the TDE mounting medium from blocking breathing through the lateral spiracles. Using an inverted confocal spinning disk microscope from Nikon using a 10x/0,45 PL APO objective and a sCMOS camera (Orca Flash4, Hamamatsu), the anterior half of each pupa was imaged every 5min, throughout a 150-250 µm z-stack with 1-2µm spacing, at 29°C.

For live imaging of neck folding dynamics, pupae were mounted and imaged as described before (Villedieu et al., 2023). For live imaging of the pre-fold pupal neck, or Toll-2::GFP localization over time, the pupae were mounted and imaged in the same way, except that images were taken at every 5 min, throughout 30 step z-stacks with 0.5µm spacing.

The images for the screening of TLR and LRR surface molecules were taken live, but at a single time point between 13h to 15h APF, throughout a z-stack of 40 µm, with 1 µm spacing, using an inverted confocal spinning disk microscope from Nikon or Carl Zeiss using a 40x/1.4 OIL DIC H/N2 PL FLUOR objective and sCMOS camera (Orca Flash4, Hamamatsu).

#### Larval and pre-pupal tissue fixation, staining and imaging

Dissected larval eye-antennal imaginal discs were fixed in 4% paraformaldehyde:PBS for 20 minutes at room temperature and washed with PBS 0.01% Tween-20 prior to DNA staining by 40,6-diamidino-2-phenylindole (DAPI), and mounted in Vectashield mounting medium.

For dissection of pre-pupal tissues, first a small excision was made at the posterior end of the pupal case, to prevent the pupa from bursting under pressure, and then the pupa was cut in half. The pre-pupal samples were fixed at the stage in 4% paraformaldehyde:PBS for 20 minutes at room temperature and washed with PBS 0.01% Tween-20. The pupal case and remainders of associated larval tissues were removed following fixing, and pre-pupal tissue was mounted in Vectashield mounting medium.

For samples older than 12h APF, pupae were dissected, fixed and stained as previously described (Villedieu et al., 2023).

All fixed samples were imaged using an inverted confocal spinning disk microscope from Nikon or Carl Zeiss using either 40x/1.4 OIL DIC H/N2 PL FLUOR, 60x/1.4 OIL PL APO or 63x/1.4 OIL DICII PL APO objectives and sCMOS camera (Orca Flash4, Hamamatsu).

### Quantitative analysis of pre-fold tissue morphogenesis

#### Image processing

Time-lapse images of ECad::GFP signal were projected with the *local z-projector* plugin of Fiji, and segmented using a cellpose segmentation model trained on images of ECad::GFP signal in dorsal head and thorax tissues. Segmented images were manually corrected prior to quantification and analysis.

#### Quantification of cell number, shape, divisions, delaminations and rearrangements and their morphogenetic contributions

Projected and segmented images were used for quantification and analysis of cell behaviors and their morphogenetic contributions as previously described (Guirao et al., 2015). Analysis was restricted to a segmented and tracked neck mask defined by *DfdGal4>LifeAct::Ruby* expression at the first time point of imaging, unless otherwise stated.

#### Quantification of AP neck thickness

For quantification of AP neck thickness over time, for each time point analyzed, number of cell centroids found along the AP axis of the neck mask, within a sliding window of mean ML cell aspect size was averaged over the ML length of neck region quantified (medial or lateral).

#### Quantification of ML tissue flows

Tissue flow speed was quantified along the ML axis as described before (Guirao et al., 2015), and averaged within each half Dfd compartment (defined by the two halves of the neck mask as separated by the tissue midline).

### Quantification of neck fold deepening dynamics

Spatial and temporal registration of time-lapse movies, tracking of apical neck fold front and quantification of neck fold depth was performed as described before (Villedieu et al., 2023), except that we automated the initial tracking of the apical fold front based on standard watershed algorithms used on topographical z-maps of tissue depth, obtained by the *local z-projector* plugin of Fiji (Schindelin et al., 2012).

For each time point *t*, mean fold deepening speed is calculated as the difference between mean fold depth at time *t* and at time *t-1*, divided by the time step separating each successive imaging. The quantifications of neck folding dynamics for conditions *Dfd>Sqh^RNAi^, Dfd>Rok^RNAi^, Dfd> Mbs^RNAi^* and *Dfd>Toll-8^RNAi^*, *Dfd>Diap1,* presented in Figure S5 and S6, we used our previously published data (Villedieu et al., 2023).

### Laser ablations

Laser ablations to measure tissue recoil velocity were performed using an inverted laser scanning microscope (LSM880 NLO, Carl Zeiss) equipped with a multi-photon Ti::Sapphire laser (Mai Tai HP DeepSee, Spectra Physics). Ablations were performed in pupae aged between 16 to 22 hAPF Ecad:: GFP and *DfdGal4>LifeAct::Ruby* and imaged in single-photon bidirectional scan mode every 1s for 30s with a 40x/1.3 OIL DICII PL APO (UV) VIS-IR (420762-9800) objective. A small 8.6×17.2µm rectangular region of interest (ROI), positioned at the neck/thorax interface, was ablated as previously described69,74,75. The initial tissue recoil velocity, which is proportional to the stress33 was determined between t=1s and t=7s following ablation.

### Image processing for display

All images and movies are maximal z-projections and were subjected to image processing for display purposes (denoising and contrast enhancement) using Fiji77, unless otherwise stated. For 3D representations of confocal and microCT images, we used the Imaris software (https://imaris.oxinst.com/).

### Statistics and reproducibility

Sample sizes vary in each experiment and reported on each figure or the corresponding figure legend. For any given genotype, animal samples were randomly selected for experimental analyses. Each experiment was repeated at least twice, and images were selected to portray representative animals. Animals that were incorrectly mounted, damaged, or that died during imaging were discarded in subsequent analysis. For plots of data over time, each error area represents the standard error to the mean (sem). For statistical comparison of two conditions in these plots, a Welch test was performed for each time point, and the associated p-value was plotted as a horizontal bar (white: p>0.05, striped: p<0.05 and plain: p<0.01) on top of each graph. For box plots of initial AP neck thickness, initial number of cells in different neck regions or recoil velocity upon laser ablation, the median, and the upper and lower quartiles are shown, as well as the individual points making up each data set. Comparisons of two distributions were performed using Welch tests. For each figure, the statistical tests used to assess significance are stated in the figure legends. Statistical analyses were performed using Matlab.

### Data availability

The image datasets generated are available from the corresponding author upon reasonable request.

---

**Supplementary Table 1.**
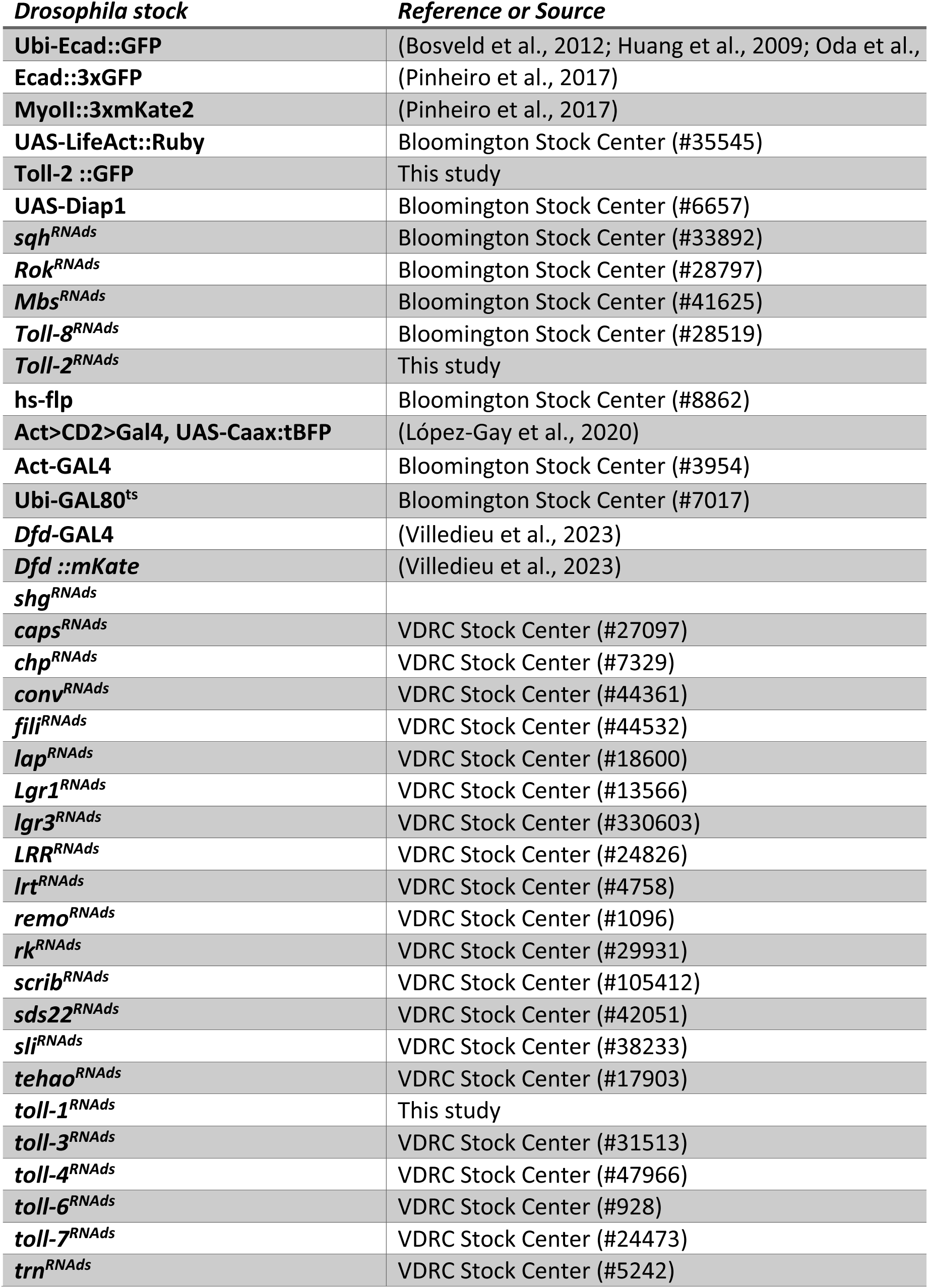
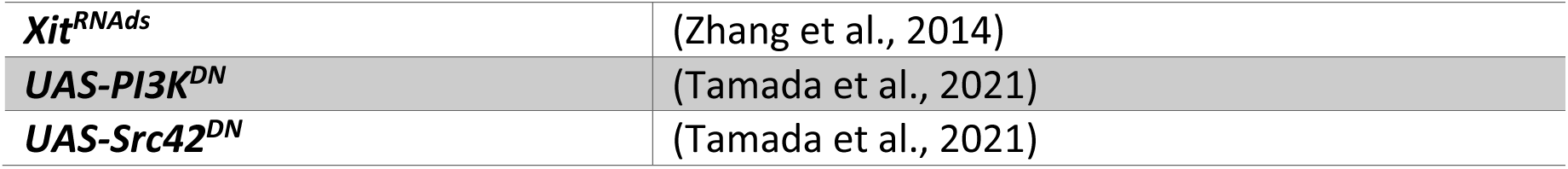
*Drosophila* alleles and transgenes.

| ***Drosophila* stock** | **Reference or Source** |
| --- | --- |
| **Ubi-Ecad::GFP** | (Bosveld et al., 2012; Huang et al., 2009; Oda et al., |
| **Ecad::3xGFP** | (Pinheiro et al., 2017) |
| **MyoII::3xmKate2** | (Pinheiro et al., 2017) |
| **UAS-LifeAct::Ruby** | Bloomington Stock Center (#35545) |
| **Toll-2 ::GFP** | This study |
| **UAS-Diap1** | Bloomington Stock Center (#6657) |
| ***sqh*<sup>RNA*ds*</sup>** | Bloomington Stock Center (#33892) |
| ***Rok*<sup>RNA*ds*</sup>** | Bloomington Stock Center (#28797) |
| ***Mbs*<sup>RNA*ds*</sup>** | Bloomington Stock Center (#41625) |
| ***Toll-8*<sup>RNA*ds*</sup>** | Bloomington Stock Center (#28519) |
| ***Toll-2*<sup>RNA*ds*</sup>** | This study |
| **hs-flp** | Bloomington Stock Center (#8862) |
| **Act>CD2>Gal4, UAS-Caax:tBFP** | (López-Gay et al., 2020) |
| **Act-GAL4** | Bloomington Stock Center (#3954) |
| **Ubi-GAL80<sup>ts</sup>** | Bloomington Stock Center (#7017) |
| ***Dfd*-GAL4** | (Villedieu et al., 2023) |
| ***Dfd* ::*mKate*** | (Villedieu et al., 2023) |
| ***shg*<sup>RNA*ds*</sup>** |  |
| ***caps*<sup>RNA*ds*</sup>** | VDRC Stock Center (#27097) |
| ***chp*<sup>RNA*ds*</sup>** | VDRC Stock Center (#7329) |
| ***conv*<sup>RNA*ds*</sup>** | VDRC Stock Center (#44361) |
| ***fili*<sup>RNA*ds*</sup>** | VDRC Stock Center (#44532) |
| ***lap*<sup>RNA*ds*</sup>** | VDRC Stock Center (#18600) |
| ***Lgr1*<sup>RNA*ds*</sup>** | VDRC Stock Center (#13566) |
| ***lgr3*<sup>RNA*ds*</sup>** | VDRC Stock Center (#330603) |
| ***LRR*<sup>RNA*ds*</sup>** | VDRC Stock Center (#24826) |
| ***Irt*<sup>RNA*ds*</sup>** | VDRC Stock Center (#4758) |
| ***remo*<sup>RNA*ds*</sup>** | VDRC Stock Center (#1096) |
| ***rk*<sup>RNA*ds*</sup>** | VDRC Stock Center (#29931) |
| ***scrib*<sup>RNA*ds*</sup>** | VDRC Stock Center (#105412) |
| ***sds22*<sup>RNA*ds*</sup>** | VDRC Stock Center (#42051) |
| ***sli*<sup>RNA*ds*</sup>** | VDRC Stock Center (#38233) |
| ***tehao*<sup>RNA*ds*</sup>** | VDRC Stock Center (#17903) |
| ***toll-1*<sup>RNA*ds*</sup>** | This study |
| ***toll-3*<sup>RNA*ds*</sup>** | VDRC Stock Center (#31513) |
| ***toll-4*<sup>RNA*ds*</sup>** | VDRC Stock Center (#47966) |
| ***toll-6*<sup>RNA*ds*</sup>** | VDRC Stock Center (#928) |
| ***toll-7*<sup>RNA*ds*</sup>** | VDRC Stock Center (#24473) |
| ***trn*<sup>RNA*ds*</sup>** | VDRC Stock Center (#5242) |
| *Xit<sup>RNAi</sup>* | (Zhang et al., 2014) |
| *UAS-PI3K<sup>DN</sup>* | (Tamada et al., 2021) |
| *UAS-Src42<sup>DN</sup>* | (Tamada et al., 2021) |

## Supplementary Figures

**Figure S1:**
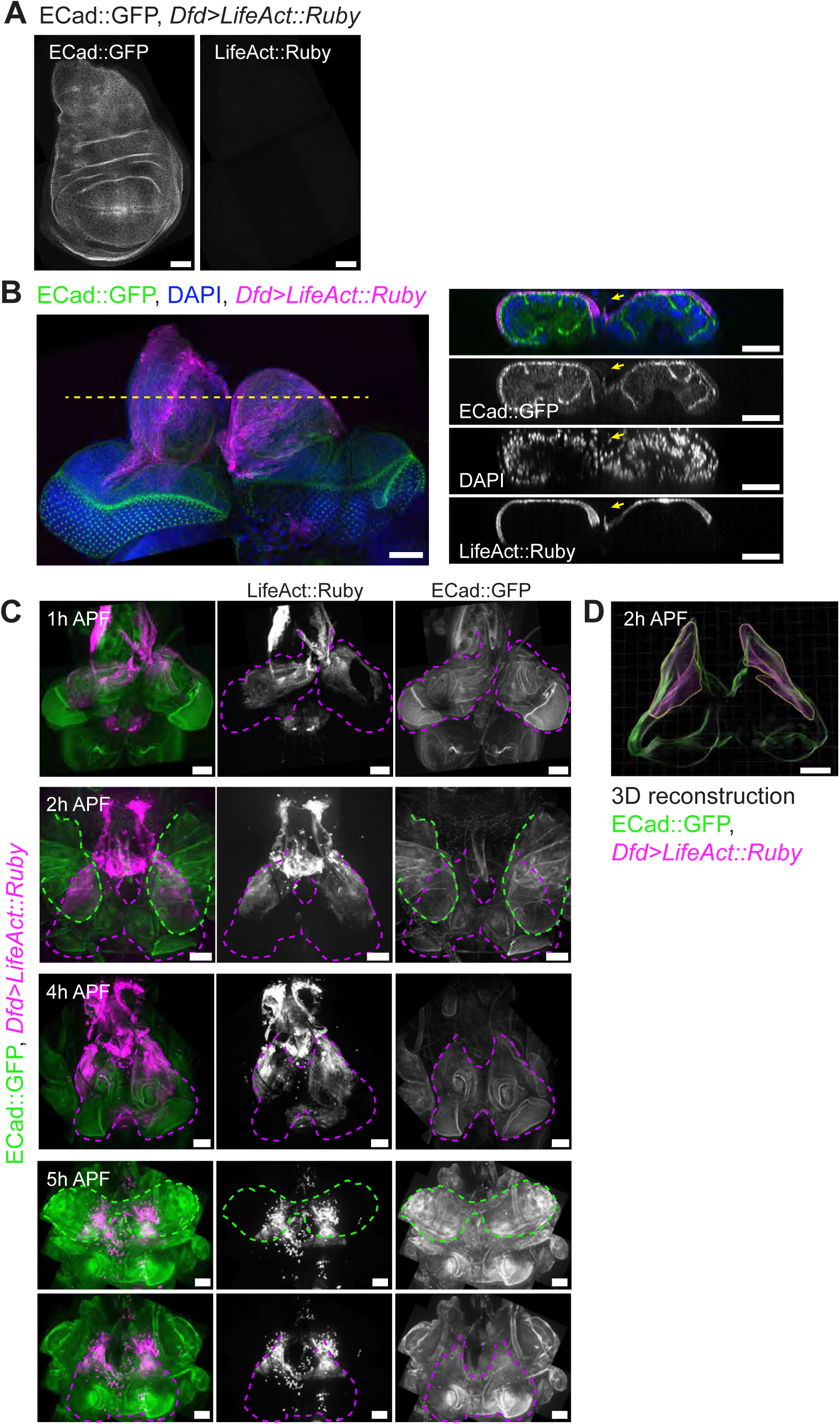
Expression of *DfdGal4>UAS-LifeAct::Ruby* during larval and pupal stages. A: 3^rd^ instar wing disc expressing *UAS-LifeAct::Ruby* under *DfdGal4* control. No *LifeAct::Ruby* expression was detected in the wing imaginal disc. Scale bars = 50 μm. B: Pair of 3^rd^ instar eye-antennal discs expressing *UAS*-*LifeAct::Ruby* under *DfdGal4* control, with the inter-antenal connection. Scale bars = 50 μm. C: Images of *DfdGal4>UAS-LifeAct::Ruby* expressing pupa, fixed at the indicated time points. Green dashed lines outline position of wing imaginal discs/nota, magenta dashed lines outline those of eye-antennal imaginal discs. For 5 hAPF, we are presenting two projections of the same image stack. Images presented at top shows projections over the entire tissue depth, while those shown at the bottom panels are projections over a lower portion of the stack, corresponding to eye-antennal discs, excluding the image slices corresponding to the closing dorsal thorax. Scale bars = 50 μm. D: 3D reconstruction of the eye-antennal discs from E-Cad::GFP and *Dfd>LifeAct::Ruby* signals at 2 hAPF, image shown in C. Eye-antennal discs represented in green, *Dfd+* compartment represented in magenta. Scale bar = 100 μm.

**Figure S2:**
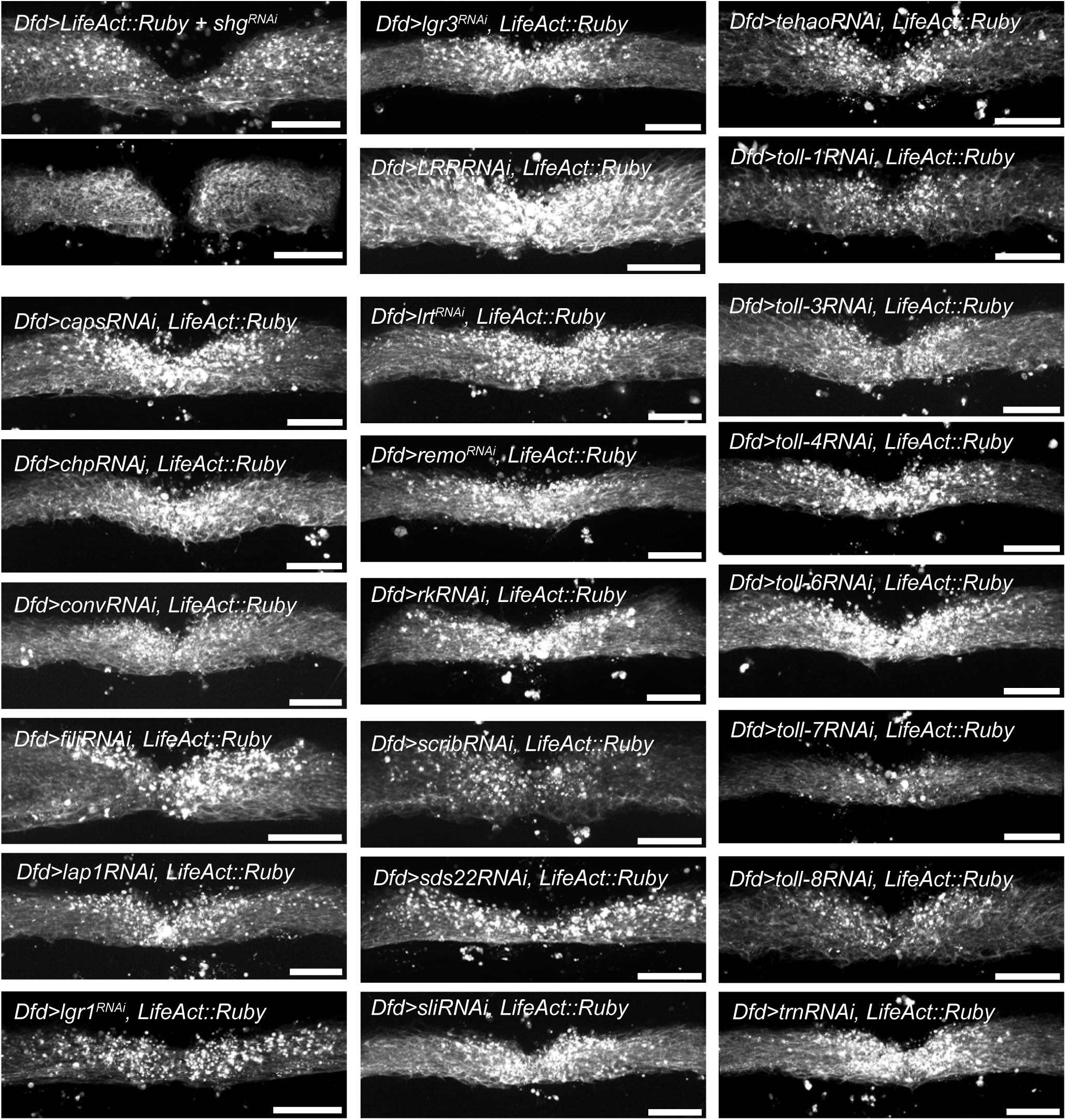
Screening for regulators of *Dfd*+ compartment shape in the pupa. Maximum signal projections of *UAS-LifeAct::Ruby* expression under *DfdGal4* control, projected along the apicobasal tissue height, in wildtype animals or in animals also expressing *RNAi* against the indicated genes. Note that for *UAS-LifeAct::Ruby + UAS-shg-RNAi* condition (top left), we are showing results from two different animals representing a milder (top) and a more severe (bottom) phenotype. Scale bars=50 μm.

**Figure S3:**
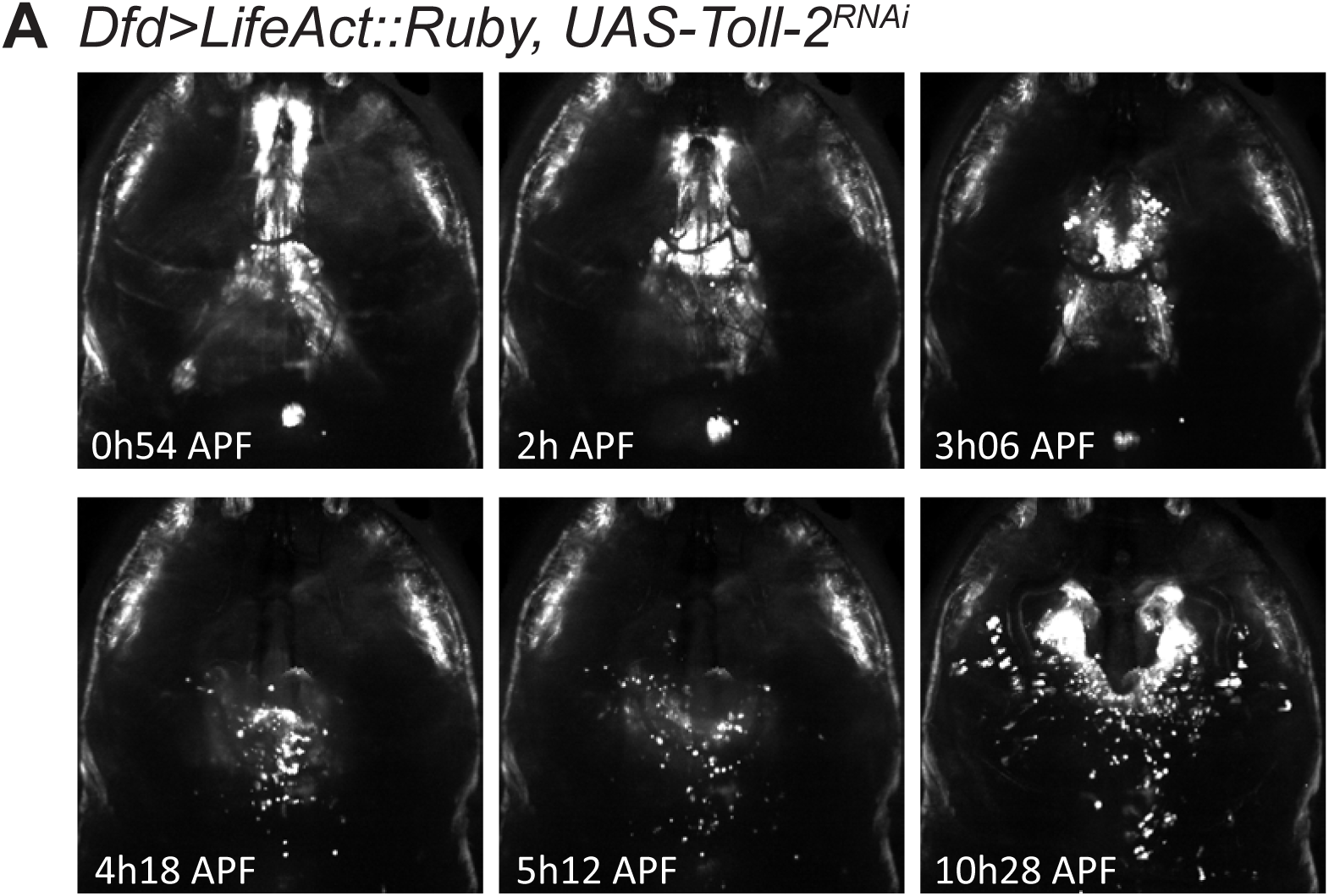
Dynamics of *Dfd*+ compartment in *Toll-2^RNAi^* pupa. Snapshots taken from Movie 3, showing position of *DfdGal4>UAS-LifeAct::Ruby* expressing tissue within *Dfd>LifeAct::Ruby + UAS-Toll-2^RNAi^*-expressing pre-pupa. Scale bars = 100 μm.

**Figure S4:**
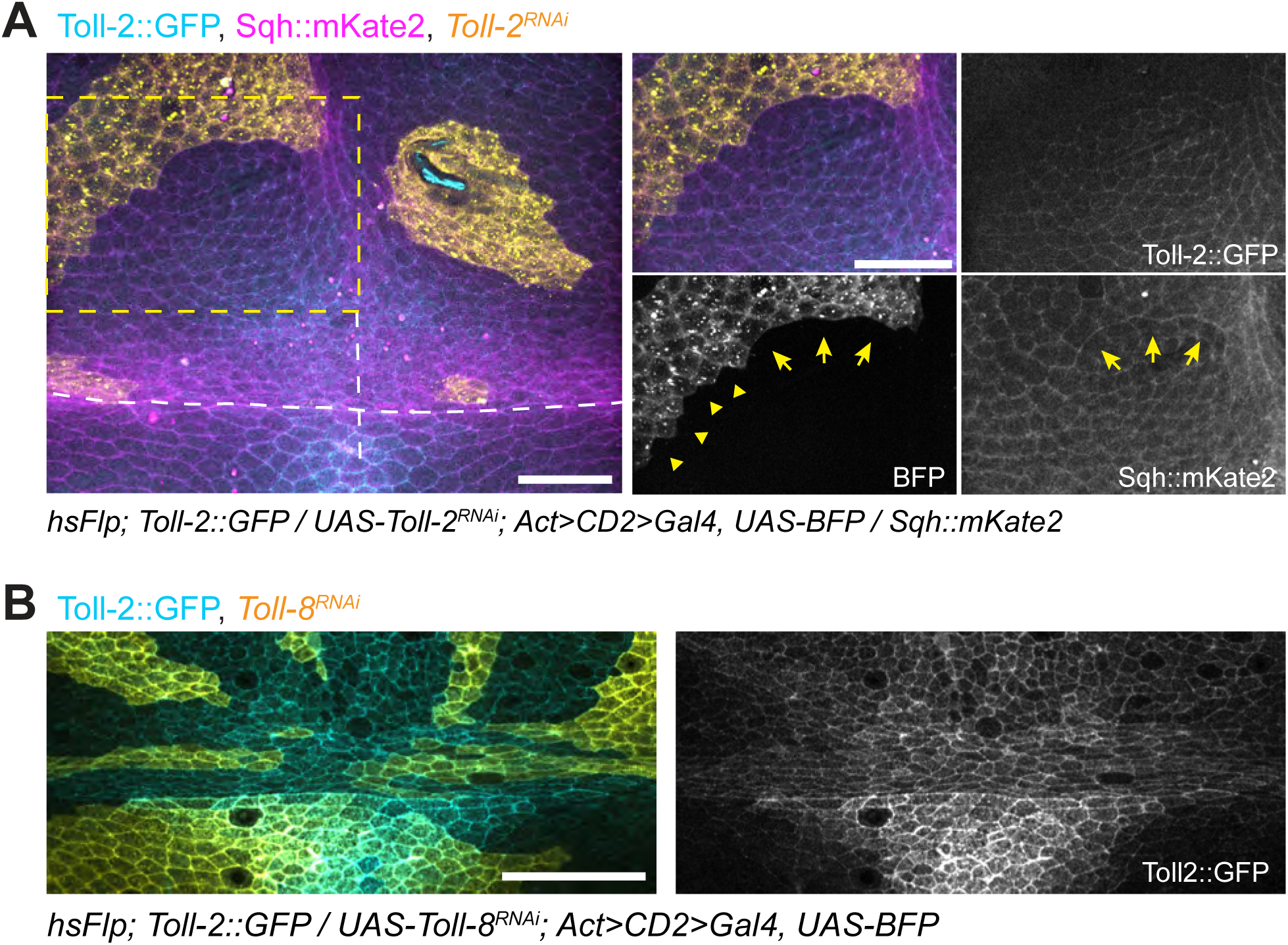
Clonal loss of Toll-2 function. A: Toll-2::GFP and Sqh::mKate2 in *UAS-Toll-2^RNAi^*-expressing head clones. White dashed lines mark midline and head/thorax boundary. Yellow dashed boxes mark region magnified in the right panels. Notice Myo II cable at smooth clone interface nearing the posterior midline (yellow arrows), and its absence on the more anterior-lateral, non-smooth interface (yellow arrowheads). Scale bars = 50 μm. B: Toll-2::GFP in *UAS-Toll-8^RNAi^*-expressing clones. No discernable change is seen in Toll-2::GFP localization in or around clone. Scale bar = 50 μm.

**Figure S5:**
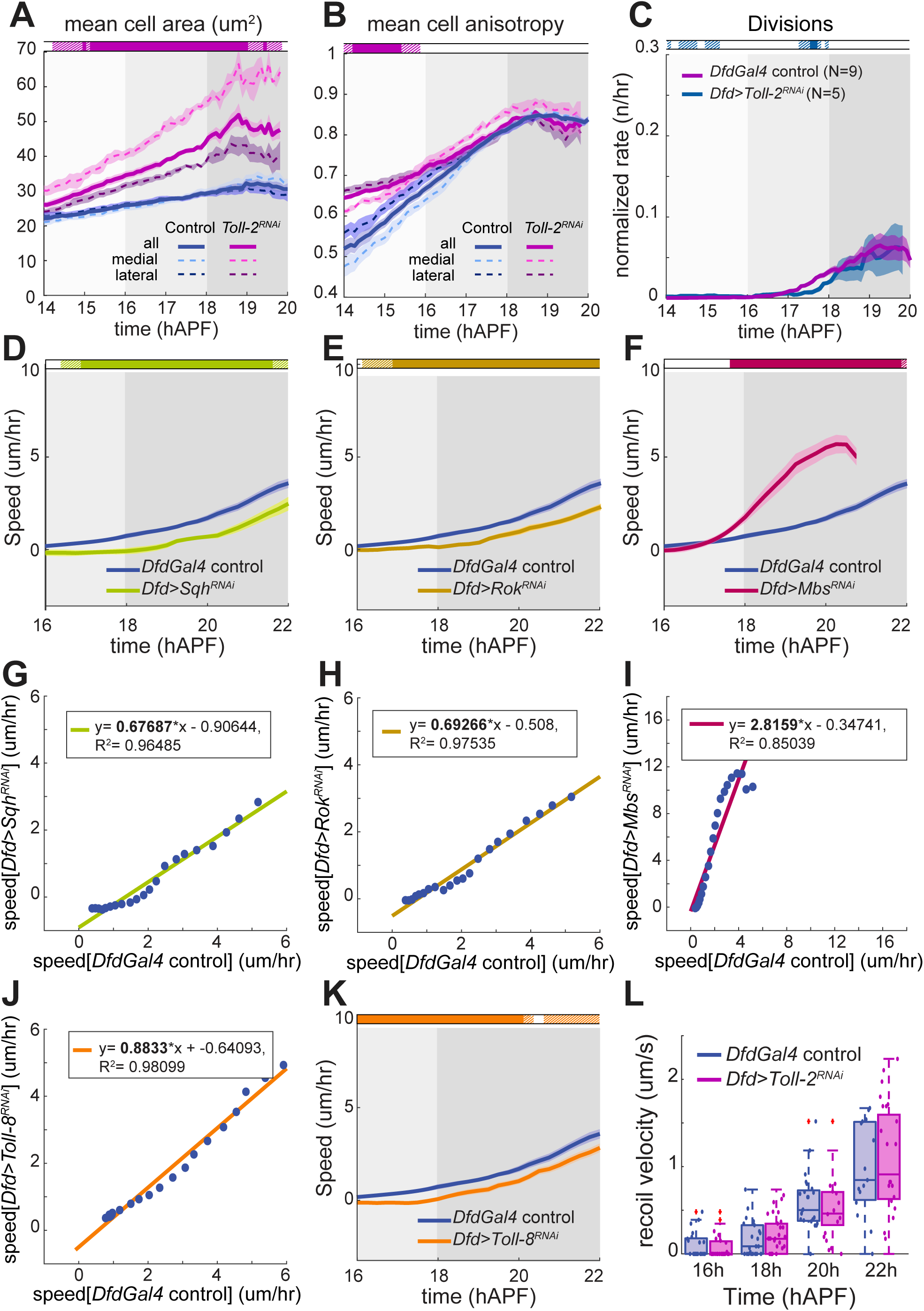
Dynamics of *Dfd+* cells during neck morphogenesis. A: Mean cell area within the *Dfd+* compartments of *DfdGal4>UAS-Toll-2^RNAi^* (magenta) and *DfdGal4>UAS-LifeAct::Ruby* control (blue) animals. Dashed lines show quantifications restricted to medial (light) and lateral (dark) neck regions as described in Figure 2. Horizontal box: p-values of Welch tests performed between control and *DfdGal4>UAS-Toll-2^RNAi^ Dfd+* compartments at successive time points (white p >0.05, striped p<0.05, solid p <0.01). *DfdGal4>UAS-LifeAct::Ruby* control N = 9; *DfdGal4>UAS-Toll-2^RNAi^* N = 5. B: Mean cell anisotropy within the *Dfd+* compartments of *DfdGal4>UAS-Toll-2^RNAi^* (magenta) and *DfdGal4>UAS-LifeAct::Ruby* control (blue) animals. Dashed lines show quantifications restricted to medial (light) and lateral (dark) neck regions as described in Figure 2. Horizontal box: p-values of Welch tests performed between control and *DfdGal4>UAS-Toll-2^RNAi^ Dfd+* compartments at successive time points (white p >0.05, striped p<0.05, solid p <0.01). *DfdGal4>UAS-LifeAct::Ruby* control N = 9; *DfdGal4>UAS-Toll-2^RNAi^* N = 5. C: Mean cell division rates within the *Dfd+* compartments of *DfdGal4> UAS-LifeAct::Ruby* control (magenta) and *DfdGal4> UAS-LifeAct::Ruby + UAS-Toll-2^RNAi^* (blue) animals, normalized to the total number of cells within analyzed region. Horizontal boxes: p-values of Welch tests performed between control and experimental conditions at successive time points (white p >0.05, striped p<0.05, solid p <0.01). N = 9 pupae. *DfdGal4>UAS-LifeAct::Ruby* control N = 9; *DfdGal4>UAS-Toll-2^RNAi^* N = 5. D: Plot showing mean speed of fold deepening over time, in *DfdGal4>UAS-sqh^RNAi^* (green) and *DfdGal4* control (blue) animals. Horizontal box: p-values of Welch tests performed between control and experimental conditions at successive time points (white p >0.05, striped p<0.05, solid p <0.01). *DfdGal4* control N = 9 pupae; *DfdGal4>UAS-sqh^RNAi^* N = 7 pupae. E: Plot showing mean speed of fold deepening over time, in *DfdGal4>UAS-rok^RNAi^* (yellow) and *DfdGal4* control (blue) animals. Horizontal box: p-values of Welch tests performed between control and experimental conditions at successive time points (white p >0.05, striped p<0.05, solid p <0.01). *DfdGal4* control N = 9 pupae; *DfdGal4>UAS-rok^RNAi^* N = 15 pupae. F: Plot showing mean speed of fold deepening over time, in *DfdGal4>UAS-mbs^RNAi^* (red) and *DfdGal4* control (blue) animals. Horizontal box: p-values of Welch tests performed between control and experimental conditions at successive time points (white p >0.05, striped p<0.05, solid p <0.01). *DfdGal4* control N = 9 pupae; *DfdGal4>UAS-mbs^RNAi^* N = 13 pupae. G: Mean speed of fold deepening in *DfdGal4>UAS-sqh^RNAi^* plotted against mean speed of fold deepening in *DfdGal4* control for each corresponding time point. *DfdGal4* control N = 9 pupae; *DfdGal4>UAS-sqh^RNAi^* N = 7 pupae. H: Mean speed of fold deepening in *DfdGal4>UAS-rok^RNAi^* plotted against mean speed of fold deepening in *DfdGal4>UAS-LifeAct::Ruby* control for each corresponding time point. *DfdGal4* control N = 9 pupae; *DfdGal4>UAS-rok^RNAi^* N = 15 pupae. I: Mean speed of fold deepening in *DfdGal4>UAS-mbs^RNAi^* plotted against mean speed of fold deepening in *DfdGal4>UAS-LifeAct::Ruby* control for each corresponding time point. *DfdGal4* control N = 9 pupae; *DfdGal4>UAS-mbs^RNAi^* N = 13 pupae. J: Mean speed of fold deepening in *DfdGal4>UAS-Toll-8^RNAi^* plotted against mean speed of fold deepening in *DfdGal4* control for each corresponding time point. *DfdGal4* control N = 9 pupae; *DfdGal4>UAS-Toll-8^RNAi^* N = 11 pupae. K: Plot showing mean speed of fold deepening over time, in *DfdGal4>UAS-Toll-8^RNAi^* (orange) and *DfdGal4>UAS-LifeAct::Ruby* control (blue) animals. Horizontal box: p-values of Welch tests performed between control and experimental conditions at successive time points (white p >0.05, striped p<0.05, solid p <0.01). *DfdGal4* control N = 9 pupae; *DfdGal4>UAS-Toll-8^RNAi^* N = 11 pupae. L: Distributions of recoil velocity following ablation at the neck/thorax interface, in *DfdGal4>UAS-Toll-2^RNAi^* (magenta) and *DfdGal4>UAS-LifeAct::Ruby* control (blue) animals, between 16h and 22h APF.

**Figure S6:**
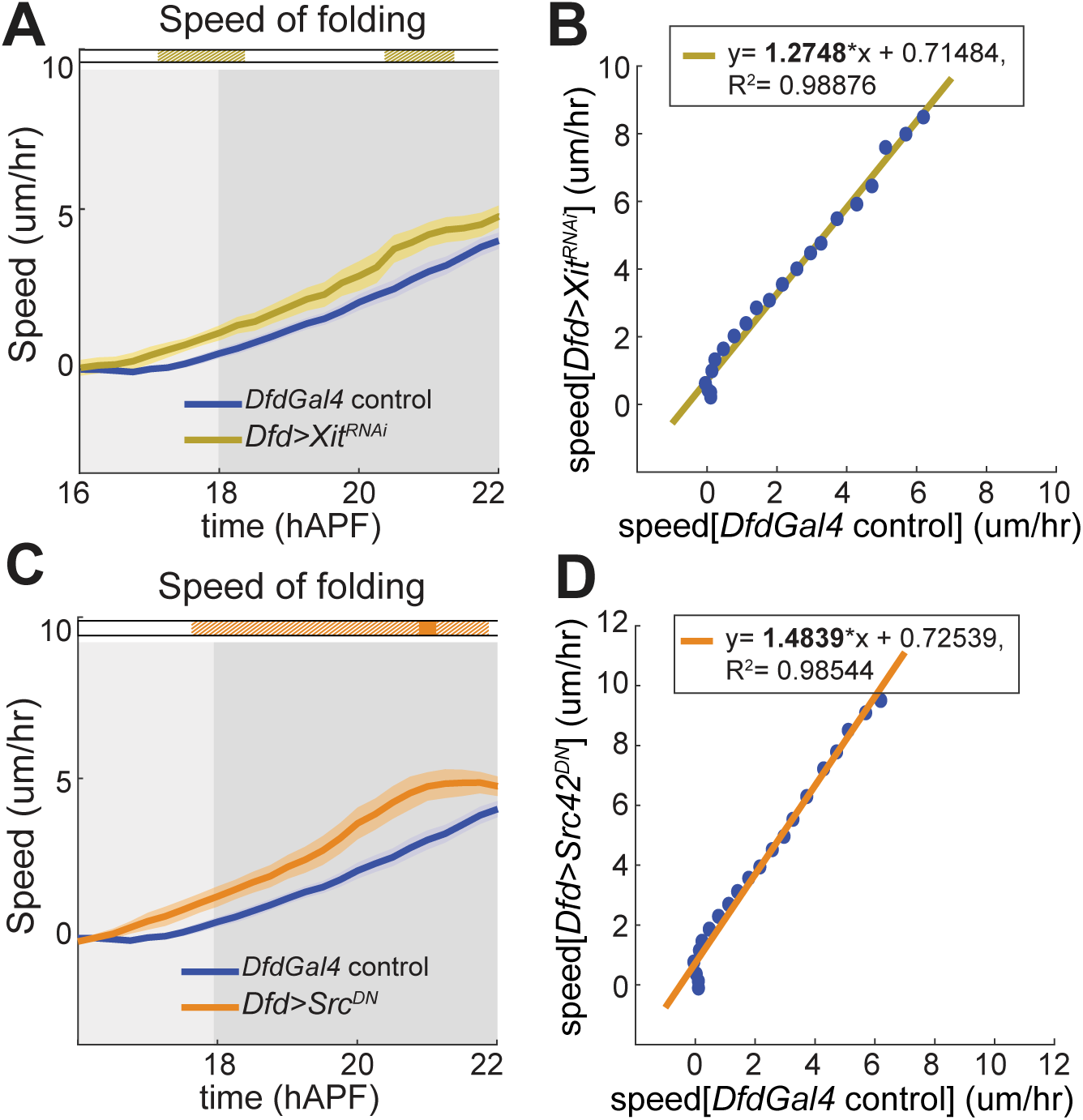
Dynamics of *Dfd*+ compartment in *Xit^RNAi^* or *Src^DN^* mutant tissues. A: Plot showing mean speed of fold deepening over time, in *DfdGal4>UAS-Xit^RNAi^* (yellow) and *DfdGal4* control (blue) animals. Horizontal box: p-values of Welch tests performed between control and experimental conditions at successive time points (white p >0.05, striped p<0.05, solid p <0.01). *DfdGal4* control N = 18 pupae; *DfdGal4> UAS-Xit^RNAi^* N = 12 pupae. B: Mean speed of fold deepening in *DfdGal4>UAS-Xit^RNAi^* plotted against mean speed of fold deepening in *DfdGal4* control for each corresponding time point. *DfdGal4* control N = 18 pupae; *DfdGal4> UAS-Xit^RNAi^* N = 12 pupae. C: Plot showing mean speed of fold deepening over time, in *DfdGal4>UAS-Src^DN^* (orange) and *DfdGal4* control (blue) animals. Horizontal box: p-values of Welch tests performed between control and experimental conditions at successive time points (white p >0.05, striped p<0.05, solid p <0.01). *DfdGal4* control N = 18 pupae; *DfdGal4> UAS-Src^DN^* N = 9 pupae. D: Mean speed of fold deepening in *DfdGal4>UAS-Src^DN^* plotted against mean speed of fold deepening in *DfdGal4* control for each corresponding time point. *DfdGal4* control N = 18 pupae; *DfdGal4> UAS-Src^DN^* N = 9 pupae. For all plots, period between onset of medial-to-lateral tissue flow and onset of folding (16h to 18h APF) shown in medium grey background; period after onset of invagination (18h APF onwards) shown in dark grey background. Reference time points are based on observed average tissue movement initiation times in controls. All quantities plotted are moving averages over a 2h sliding window, unless otherwise stated.

**Movie 1: Dynamics of *Dfd*+ compartment prior to head eversion.**

Time-lapse movie of early pupal development from 0 hAPF to head eversion, in an *E-Cad::GFP, Dfd>LifeAct::Ruby* animal. E-Cad::GFP signal in green, LifeAct::Ruby in magenta. Time-lapse images taken through the pupal case, at 29°C, with a time interval of 5 min between successive images.

**Movie 2: 3D reconstruction of eye-antennal discs at 5 hAPF.**

3D reconstructions of pupal notum (green), central nervous system (yellow) and fused eye-antennal disc (magenta) tissues, reconstructed from segmented microCT scan of a wildtype (*w; +; +*) pupa at 5 hAPF.

**Movie 3: Dynamics of *Dfd*+ compartment prior to head eversion in *Toll-2^RNAi^* pupa**

Time-lapse movie of early pupal development from 0 hAPF to head eversion, in a *Dfd>UAS-LifeAct::Ruby + UAS-Toll-2^RNAi^* animal. Time-lapse images taken through the pupal case, at 29°C, with a time interval of 5 min between successive images.

**Movie 4: Toll-2::GFP localization in the pupal neck**

Time-lapse movie of Toll-2::GFP (cyan) and Sqh::mKate2 (magenta) localization in the pupal neck, head and thorax regions, from 13 to 19 hAPF. Time-lapse images taken with a time interval of 5 min between successive images.

## Notes

### Competing Interest Statement

The authors have declared no competing interest.

